# *C*ell *3*D *P*ositioning by *O*ptical encoding *(C3PO*) and its application to spatial transcriptomics

**DOI:** 10.1101/2024.03.12.584578

**Authors:** James Cotterell, Jim Swoger, Alexandre Robert-Moreno, Heura Cardona, Marco Musy, James Sharpe

## Abstract

Current state-of-the-art spatial omics approaches suffer from the drawback that they are tissue section-based and thus inherently 2-dimensional. A full understanding of biological processes will only be possible when such data is available in 3-dimensions (3D). Here, we introduce Cell 3D Positioning by Optical encoding (C3PO) - the first technique capable of reconstructing the 3D positions of cells in a tissue, after they have been fully dissociated for single-cell omics analysis. It imposes a Cartesian coordinate system of positions on the tissue and cells of interest, before dissociation. This is created by multiple orthogonal spatial gradients of active fluorophores, carefully shaped by a 3D bleaching method, such that each position in the tissue is encoded by a unique fluorescent address. Upon dissociation of the tissue the fluorescent addresses of the cells can be read via an appropriate device (such as a FACS machine) to computationally reconstruct the tissue in 3D, before omics are performed downstream. Here, we show two proof-of-principle results for C3PO. First, pure C3PO without omics, to reconstruct the 3D geometry of a developing mouse limb bud. Second, an application of C3PO to spatial transcriptomics, revealing the expression patterns of 73 genes with interesting gene expression patterns in the developing limb.. C3PO is a genuinely novel approach to reconstruct the original 3D positions of cells in a tissue after dissociation. Combined with transcriptomics, it can play a significant role in the study of any tissue or organ in which 3D structure and geometry is important, such as developmental biology, cancer biology and neuroscience. It is not an omics technique *per se*, and in the future could be combined with the growing family of other omics technologies.

**One sentence summary:** C3PO is a novel optical technique that can preserve the 3D positional coordinates of cells after tissue dissociation, enabling a radically new approach to spatial transcriptomics.

## Introduction

Traditional approaches to molecular analysis of tissues have exhibited a trade-off. On one hand, high throughput single-cell approaches have produced comprehensive omics data (genomics, transcriptomics, metabolomics, epigenomics, proteomics) for very high numbers of cells (such as scRNASeq^1^, CyTOF^2^, CITE-Seq^3^ etc) but with no spatial information about where each cell had been located in the tissue. On the other hand, *in situ* hybridisation techniques obtain beautifully detailed spatial information (such as the Hybridization Chain Reaction, whole mount *in situ* hybridization, etc), but for only a handful of genes at a time. The new revolution of spatial omics promises to address this trade-off by providing comprehensive omics data for each cell (or region) in combination with the information of where those cells were located in the tissue^4^.

One approach to map single-cell omics into a spatial reconstruction of the organ, is to create a standardised 3D atlas of the sample, in which the expression patterns of some key marker genes are first imaged and mapped into the atlas. With enough mapped genes many positions within the tissue can be uniquely identified by specific combinations of gene expression. For example, Achim et al.^5^ mapped 98 genes into an atlas of the developing annelid *Platynereis dumerilii*, which then allowed the 3D locations of 139 single-cell transcriptomes to be mapped back into the atlas. A major drawback of this approach is the need for a prior atlas of mapped data, which requires the analysis of many samples that are believed to have the same geometric arrangement of cells or tissues (eg. a standard developmental stage of a particular developing organ). Creating standard atlases like this is a significant and time-consuming challenge^6,7^, but more importantly it is impossible to create a standard atlas for many types of 3D samples, such as tumours or mutant embryos which may have a different structure every time.

Thus the field of spatial transcriptomics has sought techniques for recording the location of cells or transcripts, without any prior knowledge/mapped data of the sample. There are currently two main approaches to achieve this^4^, which preserve spatial information in different ways: (A) Imaging based approaches preserve 2D spatial information by not dissociating the 2D section during analysis. They quantify the spatial distribution of mRNAs by directly imaging on the slide a fluorescently-labelled process - either sequential probe hybridization or *in situ* sequencing by synthesis, within the intact sections of tissue. This category includes seqFISH^8,9^, seqFISH+^10^, merFISH^11^, osmFISH^12^, RNAscope^13^, HybISS^14^, FISSEQ^15^, BaristaSeq^16^, and STARmap^17^. (B) By contrast, Next Generation Sequencing (NGS)-based approaches do dissociate the tissue section but preserve 2D spatial information by having previously encoded the 2D locations of each transcript using position-specific sequence barcodes (encoded in first-strand cDNA synthesis oligonucleotides anchored to unique known positions on the glass slide). Once first-strand cDNA synthesis has been performed, the tissue can be digested and library prep carried out on the pool of cDNAs from the entire tissue. The original spatial location of each transcript is later recovered upon sequencing of its mRNA and associated spatial barcode. This second category includes slide-seq^18^, HDST^19^, 10x Visium^20^, stereo-seq^21^ and seq-scope^22^.

An important limitation of the vast majority of the current state-of-the-art techniques (both imaging-based and NGS-based) is that they are inherently 2-dimensional - they all depend on cutting the tissue into 2D sections and analysing these on glass slides (Fig. 1a). A full understanding of tissue function requires 3-dimensional data of the relevant biological components, since only then will the full signalling environment for each cell be revealed. Quasi-3D reconstructions of a tissue can be created by combining the results of many 2D sections, but this produces alignment problems, is highly labour intensive and expensive. STARmap^23^ can perform genuine 3D transcriptomics via a hydrogel (which could also be applicable to many of the imaging based approaches) but for a limited tissue depth and only for a limited number of genes. Furthermore, the NGS-based approaches do not have a one-to-one mapping between cells and spatial barcodes and therefore some degree of deconvolution is required after the technique to elucidate which combination of cell types were present at each interrogated spatial location.

**Figure 1:**
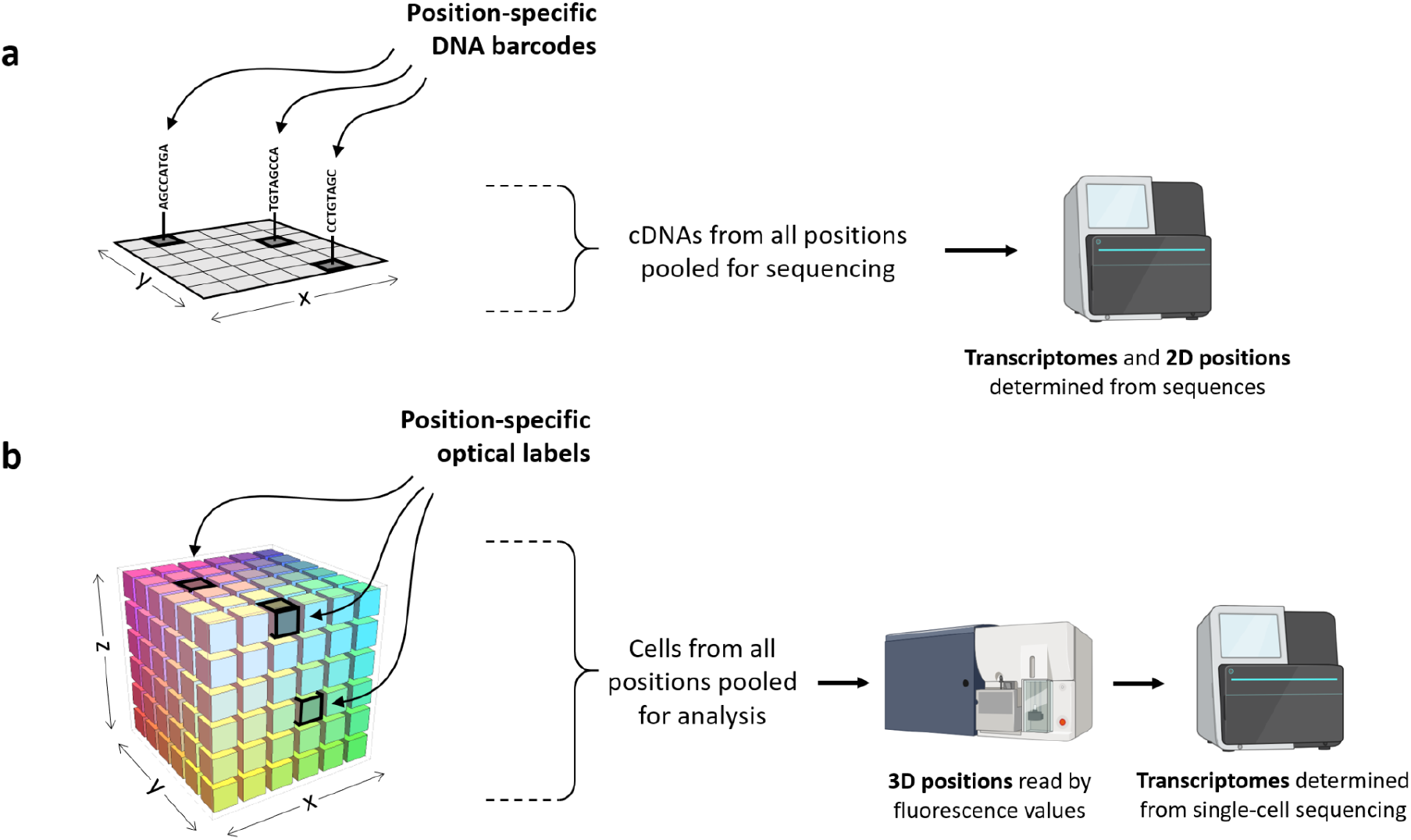
C3PO is fundamentally different to current state-of-the-art spatial omics approaches. (a) Current positional-labelling techniques are limited to 2D, as they depend on pre-locating DNA barcodes on a glass slide, which captures where each transcript was. The barcodes allow all cDNAs to be pooled for sequencing, and their original position can be recovered from the sequence. (b) C3PO optically bleaches a 3D coordinate framework of fluorescence values into the intact tissue (a different fluorescent colour for each of the 3 dimensions x, y and z). The cells can then be disassociated and pooled, and subsequently their positions recovered by recording the 4 fluorescent channels, prior to transcriptomics.

To go beyond the current state-of-the-art we sought to develop a method that could impose a comprehensive set of 3D labels onto cells before cutting or dissociating the cells for analysis - i.e. while the tissue or organ is still fully intact. We developed Cell 3D Positioning by Optical encoding (C3PO), a novel technique which intrinsically allows the preservation of 3D location information for each cell. Our new method, like the NGS approaches, also works with position-specific labels - but they are optical labels, not sequence-based labels. A simple version of optical labelling has been previously demonstrated in NICHE-Seq^24^, in which photoconversion of fluorescent proteins was used to define a distinct cell population (region of interest, ROI) within a tissue. TATTOO-Seq^25^ went further and delineated 3 ROIs within each sample, by using different degrees of photoconversion. One limitation of both these techniques is the need for a transgenic animal, which is not possible in many studies. Light-Seq can optically define ROIs in non-transgenic tissue, in this case by directing a light-dependent barcoding reaction^26^, but it is limited to 2D tissue sections. The strongest limitation of all these prior optical techniques is that they delineate just a few discrete ROIs (which must therefore be chosen by the user from the image). TATTOO-Seq was used to create an expression atlas with 14 regions, but only by combining the results of multiple limb buds^25^. None of these techniques can reconstruct the full expression patterns (nor the 3D geometry) from a single sample.

Rather than using ROIs, C3PO creates smooth gradients of fluorescence intensity through the tissue from one side to the other, by a careful bleaching protocol. By arranging three independent gradients orthogonal to each other, C3PO imposes a 3D Cartesian coordinate system across the whole tissue, so that after dissociation each cell individually carries an optical address of its original position within the tissue. In this study we describe the technique, and provide the first proof of concept that C3PO works on an organ - the embryonic mouse limb bud. We have demonstrated that the principle works in 3D to reconstruct the correct 3D geometry of the organ, and in 2D we have demonstrated its ability to perform spatial transcriptomics recording the patterns of 19,453 genes in a single experiment.

## Results

The core concept of C3PO is the creation of independent 3D spatial active fluorophore gradients within a biological sample - together these will optically encode the positions of the cells. Although a number of methods could be used to generate these spatial gradients, e.g. photoactivation/conversion, dye diffusion, or spatially controlled genetic encoding of fluorescent proteins, we chose to investigate the use of ***photobleaching*** of existing fluorophores as the most practical and broadly applicable approach. Our idea was to introduce a spatially uniform concentration of the fluorophores of interest throughout the sample (which can be done more easily than “writing” a controlled gradient in a complex 3D tissue), and then to de-activate a subset of these fluorophores by spatially controlled photobleaching. If done properly, this should result in smooth spatial gradients of the concentration of the ***active fluorophores*** in the sample. A key technical challenge was therefore to explore whether suitable gradients could be generated by the careful bleaching of different fluorophores within the tissue. This would require controlling the spatial pattern of bleaching accurately, such that tissue at one end of a given axis would be strongly bleached, while remaining unbleached at the other end with an incremental transition of bleaching in between. More specifically, all points in the tissue at the same position along this axis (i.e. all points in a plane perpendicular to the axis) should be bleached by the same amount, which would ideally define a Cartesian coordinate system. Additionally, to maximise the precision with which we can spatially map the cells we wanted to maximise the dynamic range of resulting active fluorophore gradient that is produced along the axis (unbleached to strongly bleached). Intrinsic fluorescence of the tissue sample (from fixation etc.) is weak and non-uniform, and our first task was therefore to search for a suitable cocktail of bright fluorophores to infuse uniformly across the whole tissue. Success in estimating the position of each cell would depend critically on the quantitative accuracy of creating and measuring the spatial gradients of fluorescence, and we identified 7 key requirements to achieve this:

First, although the goal is to encode 3 spatial dimensions, we must use 4 independent fluorophores. This is because even before bleaching, there will be some non-uniformity in the spatial distribution of all fluorophores. Two adjacent cells, which are in almost identical positions, may absorb the fluorescent dyes by different amounts, and therefore show quite different raw values of fluorescence signal. For this reason we need a labelling uniformity control - a fourth fluorophore which will not be bleached and can give an independent readout of how much dye each cell absorbed. The 3 spatial fluorescent channels can then be normalised with respect to this fourth channel.

Second, since the fluorescent channels will represent different spatial dimensions (plus labelling uniformity control), we must be able to manipulate (i.e. bleach) and measure each fluorophore as independently as possible. Cross-talk will lead to the bleached levels of a single fluorophore influencing the perceived cell position in more than one spatial dimension, which would make accurate 3D reconstruction non-trivial. We thus used a set of fluorophores with minimal spectral overlap (in both excitation and emission): Alexa405, 6FAM, Alexa647, and TAMRA could be bleached and read using distinct laser/Filter combinations (see spectra in Fig. 2a).

**Figure 2:**
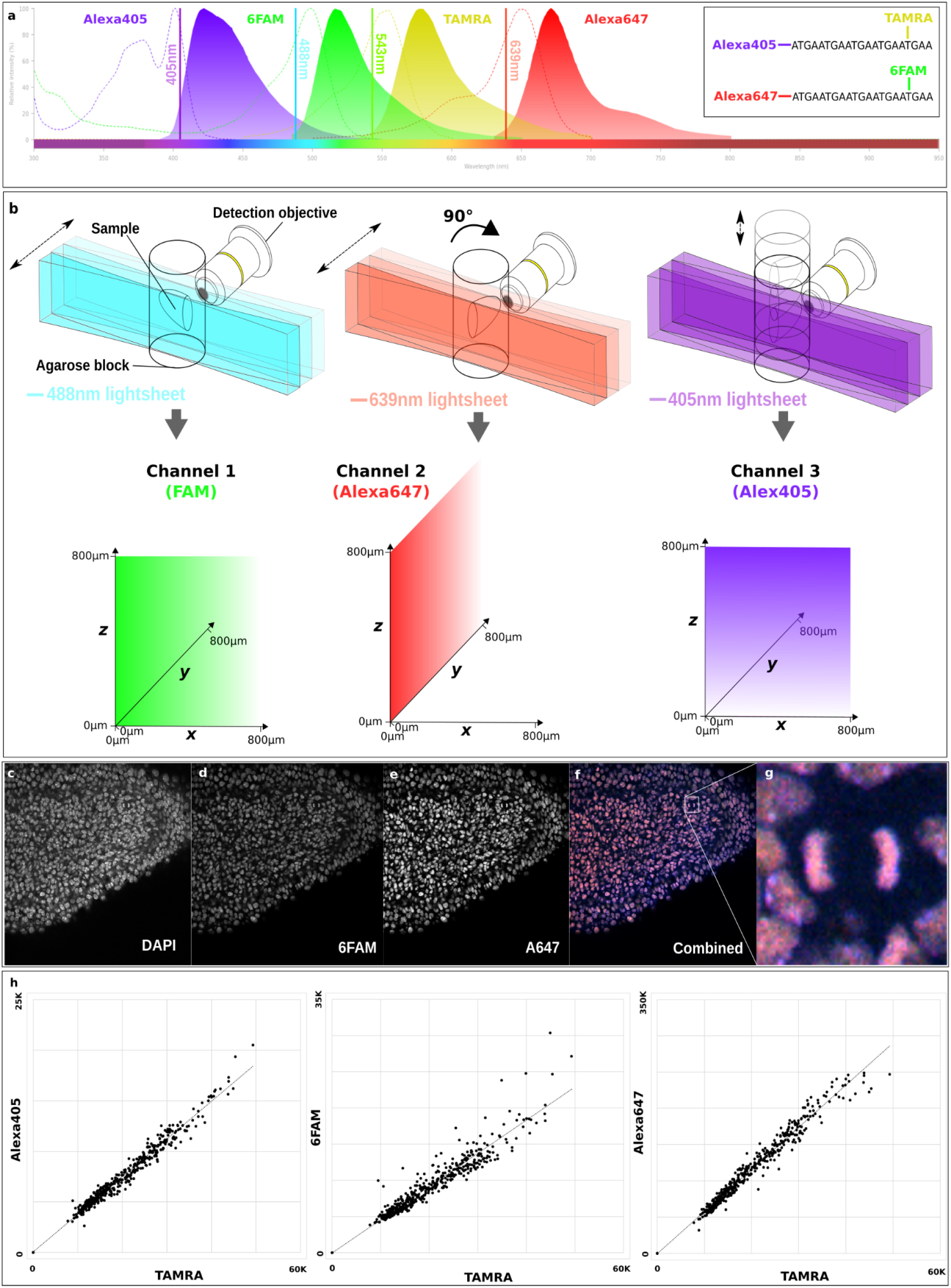
An optimal implementation of C3PO. a) Thermofisher spectraviewer illustration of the four fluorophores used for pure C3PO (Alexa405, 6FAM, TAMRA and Alexa647), three that are bleached to encode a cell’s position along each orthogonal axis and one as a labelling uniformity control. The fluorophores have minimally overlapping excitation (dashed) and emission (filled) spectra and each can be excited by a specific laser (vertical lines) with minimal excitation of the others. (inset) The two double fluorophore oligonucleotides with identical sequence used to stain the tissue stoichiometrically. b) Schematic representation of how each of the three axes is bleached (see text). For clarity we have shown the light sheet moving through the sample - in reality the sample is moved through the light sheet (c-e) Staining a tissue with only one of the oligonucleotides (6FAM/Alexa647) and DAPI and imaging tissue slabs with a confocal microscope. (f) A merged image of DAPI (blue), 6FAM (green) and Alexa647 (red) channels demonstrating colocalization of the signal. (g) A high magnification view of one cell undergoing mitosis demonstrates that the signal is localised to the chromatin during anaphase. (h) FACS analysis of a mouse limb bud stained with both oligonucleotides (unbleached) and dissociated to single cells. The signals between each pairwise combination of channels are highly correlated with Pearson’s correlation coefficients of: 0.98 for Alexa405 versus TAMRA, 0.93 for FAM versus TAMRA and 0.98 for Alexa647 versus TAMRA.

Third, in order for photobleaching to be possible at all, the sample of interest must be optically transparent. While this is naturally the case for some samples (such as the worm *C.elegans* or early-stage zebrafish embryos), for many samples of biomedical interest (such as mammalian tissues more than a few tens of microns thick) this transparency can currently only be achieved using chemical clearing techniques^27^. To be of practical use for C3PO a chemical clearing protocol must: (a) permit fluorescent labelling of the tissue, (b) render the sample sufficiently transparent to allow optical imaging and photobleaching of the desired patterns, (c) allow subsequent dissociation of individual cells for FACS analysis, and (d) be sufficiently “gentle” to not damage the features of interest in the sample (e.g. mRNA for transcriptomics studies). From multiple experiments with various clearing methods, we found that the BABB technique satisfied these requirements and was thus suitable for C3PO.

Fourth, we required a method for bleaching gradients of active fluorophore concentrations in the tissue. For this, we took advantage of Light-Sheet fluorescence Microscopy (LSFM) or Selective Plane Illumination Microscopy (SPIM)^28^, which allows precise control of the time of exposure to any given plane of the tissue. A gradient in active fluorophore concentrations along any chosen axis can be created by bleaching using a light sheet orthogonal to that axis, parking the light sheet for different amounts of time as the sample is moved through the sheet (so that one end of the sample received maximum exposure and the other end received minimal exposure, Fig. 2b left).

Fifth, we needed the spatial gradients of fluorescence to be as linear as possible. A non-linear gradient will confer greater spatial localization precision in regions where the gradient is steeper, and lower precision where the gradient is shallower. Furthermore, the use of linear monotonic gradients would allow the fluorescent ratios measured to serve as direct proxies for a cell’s estimated absolute position along each physical axis of the tissue (see results). We thus first calibrated the mapping between the SPIM’s laser power and exposure time and the level of bleaching for each fluorophore that we wished to bleach (See SM section 1). The sample was held in a fixed position relative to the light sheet and the reduction in fluorescent signal measured over time. This was performed separately for each channel (i.e. for each laser line and respective fluorophore), and this information was used to calibrate the bleaching pattern in each case.

Sixth, the three active fluorophores gradients should approximate geometric orthogonality in 3D space as closely as possible (to minimize errors associated with noise or measurement imperfections). For this, we took advantage of a SPIM microscope with the sample rotation capability^29^. After bleaching the first gradient (*x*-axis), the second monotonic gradient (the *y*-axis) could be generated by reorienting the sample by 90°, and bleaching again with the second fluorescent channel (Fig. 2b middle). The third gradient (z-axis) required a different strategy due to the geometric arrangement of the SPIM microscope. For this *z*-axis, we bleached multiple stacks through the sample with the same exposure at each slice (in other words with no spatial gradient). However, between each full stack of bleaching, we raised the sample in the vertical (*z*) direction incrementally out of the lightsheet (Fig. 2b right). Thus the top end of the sample received less bleaching than the bottom end (with an incremental gradient in between).

Seventh, since the precision of the spatial positioning depends on how well the relative signal of each bleached channel correlates with the control channel, it was important that the cells initially received the same relative amounts of all fluorophores. This is not trivial, as the dyes may be absorbed by different relative amounts for different cell types. We therefore sought a way of maximising the stoichiometry of the labelling, and chose to explore the use of oligonucleotides to covalently link multiple fluorophores to the same molecule. In order to make the chemical properties of each fluorophore as similar as possible we covalently linked fluorophore pairs (Alexa405/TAMRA and 6FAM/Alexa647) to two oligonucleotides with identical sequence (Fig. 2a inset). In order to label the tissue with these oligonucleotides we developed a specific permeabilization and binding protocol that functions in BABB cleared tissue (see methods). Confocal imaging confirmed that the fluorescence from covalently linked fluorophores emanated from the chromatin (Fig. 2c-g). Dissociation of unbleached tissue to single cells and flow cytometry analysis of all four fluorophores confirmed that although different cells absorbed significantly different amounts of the fluorescent dyes, there were very high correlations between fluorescence levels (Pearson’s coefficients from 0.93 to 0.98, Fig. 2h). In other words, if one cell absorbed 30% more dye than another, this was consistent across all 4 fluorophores - an essential criterion for C3PO to function.

These correlation graphs highlighted an important observation regarding the efficiency of labelling. The range of fluorescence intensity varied by 4-5 fold, likely reflecting natural biological variation of the cells, and differential tissue penetration. This validated the need for the fourth unbleached labelling uniformity control fluorophore (the TAMRA fluorophore, described above in point 1). If this normalization is not performed, a fluorescent signal that should determine the estimated relative position of a cell along an axis in the tissue can be confounded by non-uniformity of the labelling, resulting in a significant loss in spatial accuracy.

The value of C3PO lies in two capabilities: Firstly, C3PO must be able to reconstruct original cell positions in 3D, purely by measuring the fluorescence levels of individual cells after they have been dissociated. Given the many possible sources of quantitative error in the sample preparation and signal measurement, and the uncertainty of whether the original fluorescent levels would be maintained in cells after dissociation, it was not at all clear that this would be technically possible. Secondly, the cells must be in a suitable state for omics protocols to subsequently read out useful biological information, such as transcriptomes. Here, we demonstrate proof of concept for both of these capabilities: ‘pure’ C3PO, and the ability to perform transcriptomics afterwards. As a test sample we used the developing mouse limb bud, as it is a well-studied organ for which the expression patterns of many genes are already known (through independent conventional *in situ* labelling methods).

A couple of previous techniques have employed photoconversion (of transgenically encoded fluorescent proteins) to define up to 3 discrete ROIs in a tissue (eg.^25^). This did not provide detailed spatial information about cellular location, but rather it divided all cells into 3 bins corresponding to 3 gross regions of the tissue. The use of a single fluorescent channel also limits the pattern of ROIs that can be defined - only a 2D arrangement can be projected into the tissue. A key concept of C3PO is that by using 4 different fluorophores we should be able to project bleaching patterns in multiple different directions, and use a combinatorial approach of fluorescence ratios to define a genuinely 3D set of positions. A first test therefore (SM section S2) was to prove that 8 different ROIs could be defined in 3D by the intersection of 3 orthogonal binary ON-OFF step-patterns bleached into an E10.5 mouse limb bud. This effectively created a cube of 2×2×2 = 8 different fluorescent combinations, each corresponding to a geometric octant of the physical space containing the limb bud. Bleaching, dissociation and FACS analysis of this limb resulted in the identification of 8 populations with the expected distinct fluorescent combinations (Fig. S2). These first 3D results encouraged us to attempt a reconstruction of the full 3D geometry of the limb bud, which would require resolving many more distinct colours along each axis.

Fundamentally C3PO aims not to define ROIs (regions of interest), but to estimate the 3D position of each cell, purely from the fluorescent intensities of four fluorophores. If the positions of all the cells from a dissociated organ were correctly estimated and plotted in 3D, this would produce an estimate of the 3D shape of the tissue - a type of 3D reconstruction. The positional estimate for each cell will unavoidably have some error (due to the multiple possible sources of fluorescence variation during the course of the protocol). The larger the error, the more the reconstruction will appear as a diffuse cloud of points. A suitable test for the spatial precision of C3PO therefore requires reconstruction of a tissue with a specific shape, which is already known independently. Gene expression patterns could be used to delineate non-trivial 3D shapes, but we first wished to test pure C3PO (cell positioning) without the added technical challenges of transcriptomics. In this context, the limb bud was a useful test sample, as it includes a specific cell layer - the ectoderm - which has a thin concave shape which can be easily distinguished from a diffuse cloud of points. Since the ectoderm can be manually dissected away from the underlying mesenchyme, it provides a convenient test for whether C3PO can reconstruct a specific non-trivial geometry, without requiring the additional steps of gene expression analysis.

To this end we bleached fluorescent intensity gradients along 3 orthogonal tissue axes of a mouse limb bud (Fig. 3a), and then manually separated the ectoderm and mesenchymal tissues before dissociating the cells. To create a C3PO reconstruction of the organ the cells were passed through a FACS analyser, and the signal levels for each bleached channel were normalised with respect to the non-bleached control channel. To visualise the relative arrangement of cells in 3D space we simply plotted all the cells using their normalised fluorescence levels (Fig. 3b). We adjusted the scales of the 3 axes to match the approximate proportions of the limb bud (More physically exact scaling is possible using the predissociation lightsheet image - see SM section S7 for more details). To visualise the error in the precision of the predicted relative position of each cell, the plot of ectoderm cells (blue dots) is more informative than the mesenchymal cells (red dots). The true distribution of ectoderm cells can be seen by cutting a virtual slice through the light-sheet image (Fig. 3e), as a thin single-cell layer with a large hollow gap in the middle (blue line). Computationally extracting a similar slice through the C3PO plot of ectodermal cells (Fig. 3c,f) revealed that remarkably, the spatial distribution of predicted cell positions recreated the basic form of the true ectodermal shape - very few cells are misplaced in the centre of the limb bud, and the closed, oval ring arrangement can clearly be seen. The observed positional errors (divergence from the true shape shown in Fig. 3e) strongly suggests, for example, that dorsal cells and ventral cells are clearly separated from each other. This strong evidence of C3PO’s ability to reconstruct the shape of an organ was also repeated for two other limb buds (Fig. 3g, SM section S3 and Fig. S3). Other features of the plot also suggest a successful reconstruction: (1) fitting a curved surface through the regions of highest ectodermal density recreates an approximation of the limb bud shape with an internal concavity. Thus, the reconstruction closely reflects the known tissue configuration where the ectoderm forms a layer covering the underlying mesenchymal mass (Fig. 3d and Movie S1), (2) there are no ectodermal cells on the side of the organ where it was cut from the flank, (3) the mesenchymal core is correctly surrounded by the ectodermal cells, and (4) the plot of ectodermal cells contains a strip of higher density in the correct position of a well-known thickened region, called the Apical Ectodermal Ridge (AER). Together this data provides the first proof that the 3D distribution of cells from an organ can be reconstructed by C3PO using only the fluorescent signal from each cell after they have all been fully dissociated from the original sample.

**Figure 3:**
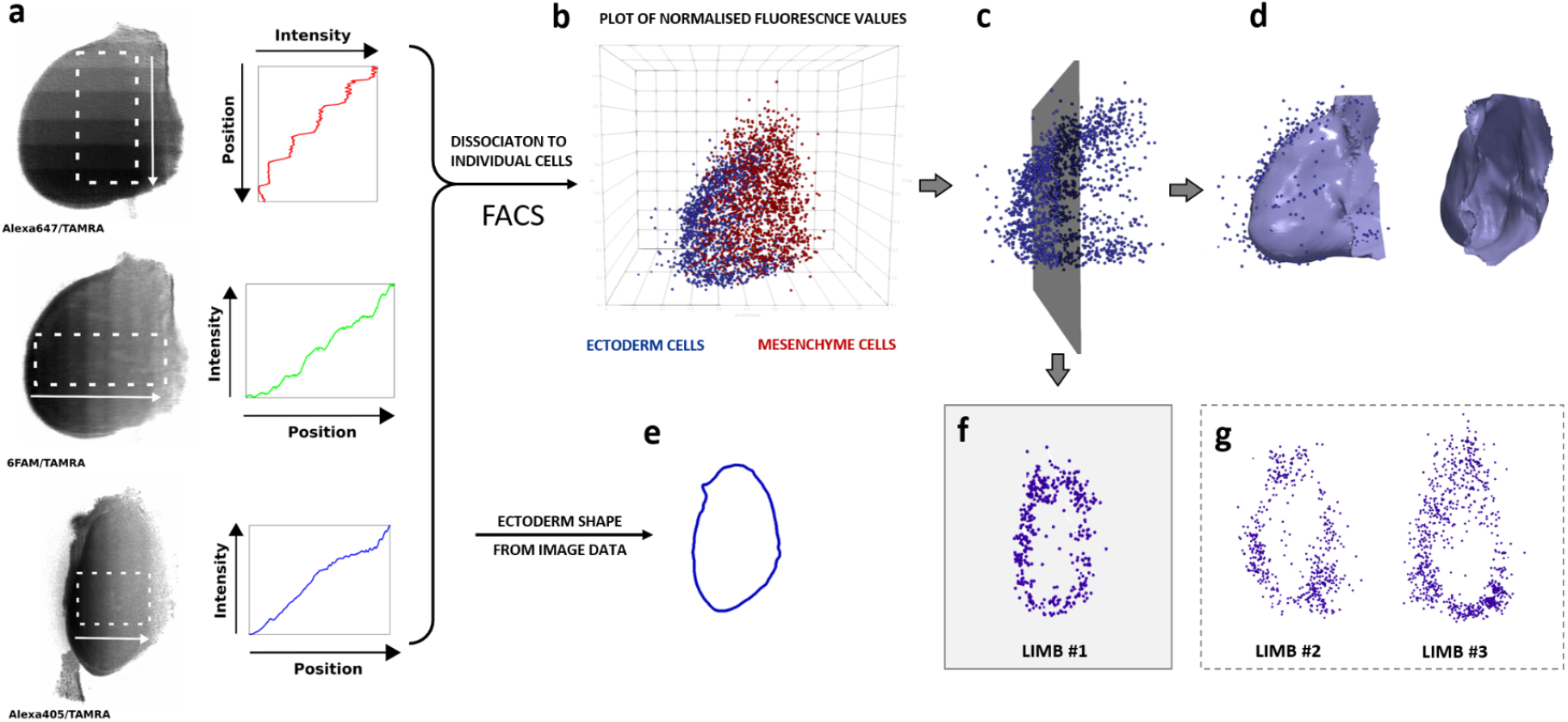
Proof-of-principle of pure C3PO. (a) Monotonic gradient bleaching of a mouse limb bud along 3 orthogonal axes. For technical convenience the gradients were bleached as step-functions with 8 increments. Sum-projection images show the respective bleached channels normalised to the labelling uniformity control channel. After bleaching the overlying ectoderm was physically dissected away from the mesenchyme and the tissues dissociated separately to single cells. The individual cells were then run through FACS analysis and the fluorescent signals in the different channels recorded. (b) A 3D scatter plot of 3,133 cells, in which the position of each cell was taken from the normalised fluorescence values of the three bleached channels (ectodermal cells are blue, and mesenchymal cells are red). (c) The distribution of ectodermal cells reveals the specific shape of the limb bud. (d) A curved surface fitted through the higher density regions of the ectodermal plot recreates an approximation of the limb bud shape. (e) Virtual section through the 3D light-sheet image extracts the true shape of the single-cell ectodermal layer. (f) The C3PO reconstruction of ectodermal cell positions for a section similar to the section in (e). (g) Virtual sections through reconstructions of two other limb buds.

Having demonstrated proof of concept for pure C3PO (with no omics), we next needed to test that this experimental pipeline (labelling, clearing, bleaching, imaging, dissociation and FACS analysis) was compatible with omics, in particular to perform spatial transcriptomics. Two modes can exist for C3PO transcriptomics: (1) In *single-cell mode*, we would generate a separate transcriptome for every single cell that is FACS-analysed. This would require capturing every cell into a separate well for its own bar-code specific cDNA library synthesis (like, for example, SMART-Seq). (2) Instead, here we employed the more economical *grid-mode*. Each C3PO-analysed cell has its own predicted position in space, but we can choose to virtually divide the sample up into a grid of blocks (Fig. 4a), and then bin all the cells which fall within the boundaries of the same block. A suitable FACS machine can be programmed to sort cells into bins of choice, thereby collecting, say, 100 cells per bin, which also increases the signal-to-noise ratio compared to single-cell sequencing. An attractive feature of C3PO is that the planning of the spatial layout of blocks (the density of the grid) is independent of the bleaching procedure. Bleaching provides global coordinates, while the choice of the size and position of binned regions can be decided and implemented afterwards (unlike previous ROI-based methods).

**Figure 4:**
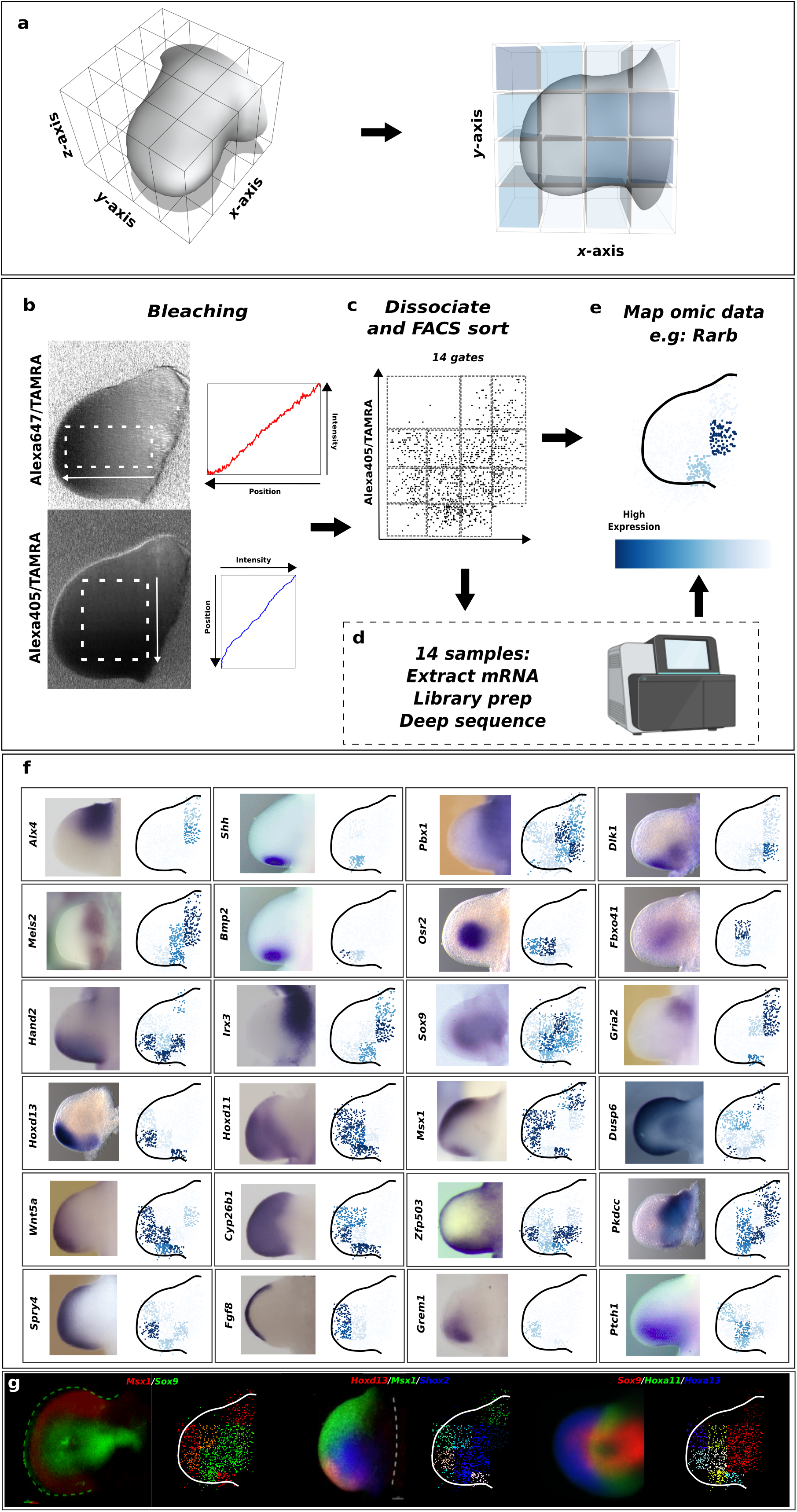
Spatial transcriptomics with C3PO in 2D-grid mode. (a) In grid mode, after bleaching and tissue dissociation cells are sorted into an organized grid of blocks that correspond to particular spatial regions of the sample (left). Omics can then be carried out on pools of cells from each of those blocks and the resulting omic data mapped back into the grid (right). Blue intensity indicates the expression level of a gene. (b) Monotonic gradient bleaching of the 2 principal axes of the mouse limb bud: proximal-distal (top) and anterior-posterior (bottom). Sum-projection images showing the respective bleached channel normalized to the labelling uniformity control channel. Plots of the signal intensities over the spaces corresponding to the dashed boxes for the two channels are shown next to the images. (c) After bleaching the tissue was dissociated and the individual cells were run through a FACS machine. Cells were ratio sorted into 14 different tubes according to their bleached channel signals normalized by the labelling uniformity control channel. The 14 regions were defined so that they encompassed the entire tissue. (d) RNAseq was carried out on the 14 individual samples. Image from Biorender. (e) Expression data could then be mapped back onto the relative position of the cells as directly measured by FACS normalized signals (i.e. the C3PO reconstruction). (f) Examples of C3PO enabled spatial transcriptomics of 24 key genes involved in mouse limb bud development. Positions of all cells are plotted individually, while the gene expression intensity is the average for all cells within a given block (cf. panel c). The C3PO reconstruction is shown next to the known gene expression profile as measured by in situ hybridization. Expression level is indicated by the depth of blue shading. Whole mount *in situ* hybridisation (WMISH) patterns for *Osr2, Dlk1, Fbxo41, Pkdcc*, Hoxd13 reprinted from^30^ with permission from Elsevier. *Hand2, Alx4, Irx3, Hoxd11, Grem1, Cyp26b1, Fgf8* reprinted from^31^ under a Creative Commons CC-BY-NC-ND licence, *Wnt5a, Spry4, Gria2, Pbx1* reprinted from^32^ with a licence from Copyright Clearance Center on behalf of The Company of Biologists Ltd., *Shh, Bmp2, Ptch1* reprinted from^33^ under a Creative Commons CC-BY licence, *Msx1, Meis2* reprinted from the Embrys database^34^ with permission from Hiroshi Asahara. *Sox9* reprinted from^35^ under a Creative Commons CC-BY licence, *Dusp6* generated by the Sharpe laboratory, and *Zfp503* reprinted from^36^ with a licence from Copyright Clearance Center on behalf of John Wiley & Sons - Books. (g) Examples of C3PO enabled spatial transcriptomics with multiple genes mapped simultaneously in the indicated colours. The C3PO reconstruction is shown next to the known gene expression profile as measured by multi-gene Hybridization Chain Reaction (HCR). Left and middle expression profiles reprinted from^37^ with permission from Elsevier (left Msx1/Sox9 profile was generated by combining two separate images). See HCR methods for right expression profile.

Our first test was a 1D grid mode transcriptomics along the proximal-distal axis of the mouse limb bud. The vast majority of gene expression patterns reconstructed closely resemble the known gene expression patterns from *in situ* hybridisations (See SM section S4 and Fig. S4). However, gene expression patterns in the limb have historically mostly been studied in 2D, so our most rigorous test of C3PO was to attempt 2D grid mode of the limb bud (Fig. 4a). We bleached orthogonal, incremental gradients along the two principal axes of the developing limb bud: the proximal-distal axis and the anterior-posterior axis (Fig. 4b). After bleaching, the limb tissue was dissociated enzymatically to single cells and the cells ran through a FACS sorter. We implemented 2D grid mode by creating rectangle sorting gates on a double ratio plot (Alexa647/TAMRA versus Alexa405/TAMRA), meaning that cells were normalized for the labelling uniformity control on-the-fly during sorting (Fig. 4c). We then sorted cells into the gates in serial fashion, sorting into 4 eppendorf tubes at a time consecutively until all gates had been sorted and cells exhausted. We then performed RNASeq on the 14 samples corresponding to the 14 grid regions that we ratio sorted (Fig. 4d). The measured RNASeq data from the 14 grid regions can be mapped directly back onto the C3PO tissue reconstruction (and the scatter plot from normalized FACS signal data scaled to fit the predissociation limb outline) as shown in Fig. 4e.

The results are shown in Fig. 4f - the predicted spatial patterns of 24 key genes involved in limb development, alongside images of those same gene expression patterns assessed by conventional whole-mount *in situ* hybridisation (WMISH). The C3PO predictions are remarkably similar to the WMISH results, providing strong proof of concept of this novel technique. To more extensively assess C3PO in an unbiased fashion, we analysed the results from two similarly-aged limb buds (limbs #4 and #5 - for limb #5 individual results see SM section S5 and Fig. S5). The 100 most expressed genes from each limb bud (from the RNA-Seq results) were added to the 24 genes initially analysed, resulting in a total of 138 genes (due to overlaps between the lists). We then searched for previously-published expression patterns for all these genes, in two public databases: the MGI GXD database (Jackson Laboratory), and the embrys database (Tokyo Medical and Dental University). 87 of these genes had a discernible gene expression pattern, of which 73 (=84%) showed a good or reasonable match with the prediction by C3PO (SM Fig. S6). The C3PO results of all genes detected in a single sample (a single limb) are of course captured in the same grid space, so we can visualise multiple genes at the same time with different colors (Fig. 4g). This is equivalent to fluorescent WMISH techniques (such as HCR, Hybridisation Chain Reaction) which can experimentally query the expression of 2 or 3 genes in the same sample, and are shown next to each C3PO result (Fig. 4g). In summary, we have demonstrated that the core concept of C3PO - recovering the spatial position of cells from multiple orthogonal bleached fluorescent gradients - is compatible with transcriptomics, and thereby capable of reconstructing the majority of known gene expression patterns.

Finally, we wished to determine the precision for which we could spatially localize cells using C3PO in our current optical configuration. Since we maximise the intensity range of each spatial gradient to the depth of the tissue along that axis, there is no universal absolute precision for all samples, but rather it depends on the size of the tissue being processed (the smaller the tissue the greater the precision). Calculating absolute physical precision requires converting FACS colour space to physical space. The measured fluorescence levels serve as a direct proxy for absolute physical position of the cells because we utilized orthogonal linear gradients. The relative address can be converted to a physical address by independently scaling each dimension of the FACS measured 3D fluorescent level point cloud so that it fits the physical size of each axis measured from the predissociation 3D SPIM image (See SM section S7 and table S1). Precision calculations can be found in SM section S8. Using this absolute physical measurements the precision for C3PO in the mouse limb bud can be defined as 51µm for the Alexa647 channel, 48µm for the Alexa405 channel and 52µm for the FAM channel (assuming Alexa647 and 6FAM channels used for the Proximal-Distal and Anterior-Posterior axes and Alexa405 channel used for the Dorsal-Ventral axis). However, a more appropriate description of the precision of C3PO is the number of resolvable segments per axis (NRSPA) since this is tissue-size independent (See SM section S8). NRSPA can be calculated as 14.3 for the Alexa647 channel, 8.8 for the Alexa405 channel and 14.8 for the 6FAM channel.

## Discussion

Here we introduce C3PO, a fluorescence-based approach to recover the 3D positions of individual cells after they have been dissociated from the tissue of interest. C3PO is not an omics technique *per se*, but is most usefully employed when combined with a technology such as RNA-Seq. It can perform spatial transcriptomics, but with a radically novel approach compared to current state-of-the-art techniques that are based on 2D tissue sections. By contrast, C3PO imposes an orthogonal optical-based Cartesian coordinate system on the tissue of interest. This is done using spectrally distinct fluorophores whose active concentrations follow monotonic spatial gradients that uniquely encode the positions of cells in three dimensions. Once dissociated, the fluorophore intensities of each cell can be read via an appropriate device such as FACS, microplate reader or microfluidic device in order to read the cell’s relative ‘address’ in the tissue. The measured fluorescence levels reveal the estimated relative cell position along each physical axis of the tissue, because the optical encoding comprises orthogonal monotonic gradients. If any downstream omics are then performed on the dissociated cells that omics information can be mapped back into the physical space of the sample.

We have demonstrated proof-of-principle of pure C3PO in which the relative spatial locations of thousands of cells are well-estimated, purely by reading out 4 fluorescent intensities from each individual cell after tissue dissociation. A 3D plot of these cells successfully reconstructs known geometrical features of tissue when applied to a mouse limb bud. It is striking that the quantitative values of fluorescence intensities for 3 different spatial axes, and the internal non-bleached control, are preserved accurately enough after tissue dissociation, through the whole C3PO processing pipeline, to create a good estimate of each cell’s position in space.

Beyond pure C3PO reconstruction, we also demonstrate that the concept and pipeline of C3PO is compatible with transcriptomics, and that we can use this approach to create maps of gene expression for thousands of genes. Comparison of predicted expression patterns with independent data from *in situ* hybridisation confirms that the majority of patterns match well. Furthermore, fluorescent *in situ* techniques (such as HCR), which allow multiple gene expression patterns to be mapped simultaneously, are informative since they allow the precision of the C3PO reconstruction to be explored (relative location of adjacent patterns with one sample) independently from its accuracy (reproducibility between samples). Thus far we provide proof-of-concept in grid-mode, however it is important to note that the spatial resolution of the grid can be improved in the future, and ultimately single-cell mode can also be performed in which the 3D location, and an individual transcriptome, can be determined for each cell. Future improvements to the technique are discussed below.

The principal advantage of C3PO over the slide-based spatial omics approaches is that C3PO is fully 3-dimensional whilst the slide-based approaches are only 2-dimensional. In a clinical setting such information may be essential in cancer biology for stratification of tumour type and selection of appropriate treatment regimes. Whilst, in a basic biology setting, a full understanding of tissue function will require having 3-dimensional quantitative data of the relevant biological components since only then will the full signalling environment for each cell be revealed. A further advantage of C3PO over many other spatial transcriptomic techniques is that no complex mapping or deconvolution is required^38^, since there is a straightforward one-to-one mapping between relative positional data of the cells (fluorophore intensities read via FACS machine after tissue dissociation) and omics data (performed on the sorted cells after FACS). For many of the NGS-based approaches the spots, nanoballs or beads used to anchor the spatial barcodes do not have a simple one-to-one relationship with each cell. For example, a Visium spot will typically capture the RNA from several cells and partial cells. Hence, a deconvolution algorithm needs to be applied to elucidate the cell types that correspond to each spot. Finally, C3PO should allow a significantly better capture depth than that of section-based NGS techniques (only 20k UMIs per spot quoted for 10x Visium). This is because first strand synthesis in C3PO takes place in solution in a tube or well instead of relying on RNA to diffuse from a tissue-section to the anchored barcode on an underlying slide. The thermodynamics of first-strand synthesis should therefore prove to be entropically more favourable when performing transcriptomics with C3PO than with slide-based approaches.

One potential improvement of C3PO involves mapping cells back onto the pre-dissociation 3D tissue image using a look-up table mapping colour-space in the FACS machine to colour space in the SPIM microscope (using single fluorophore labelled calibration samples). The resulting mapping using this approach maintains the original predissociation tissue geometry and cells are placed in the original predissociation image by assigning them to the pixel(s) that best match their measured colour. This approach has the advantage that even an individual cell can be mapped back to its original position in the tissue (a representative ensemble of cells is not required) and the absolute positional error could be potentially reduced. Furthermore, such an approach is not limited to a Cartesian coordinate system of orthogonal monotonic linear gradients as we have demonstrated in the current manuscript - other more complex optically encoded spatial maps could be employed (that we term ‘Digital C3PO’)

Improvements at every stage of the protocol promise to improve the accuracy of the C3PO reconstruction. Better accuracy will result from a greater dynamic range (S_range_) of signal in the bleached channels after bleaching (see SM section 8c). Greater dynamic range can be achieved by increasing S_max_ with improved labelling and brighter fluorophores for example. In addition S_min_ can be reduced by using different fluorophores with greater ‘bleachability’, or changes in the optical system to increase the amount of light reaching the sample (More powerful lasers for example). Reducing the noise in the fluorescent signal measured and bleached into the fluorophores will also increase the accuracy of the spatial reconstruction. This can be achieved with more stable laser power and more sensitive cameras for example. Using more channels to introduce more optical spatial information into the tissue should also reduce this noise. This could be achieved using multiple monotonic gradients in fluorophore signal along each of the tissue axes in parallel and integrating the signals from the different channels (this could also be done with digital C3PO but with more complex optical spatial patterns). Bleaching a more smooth gradient (more spatial increments) than used here will also improve spatial accuracy. Any improvement in the system used to read the fluorescent signals post tissue dissociation (currently FACS) to improve the signal-to-noise ratio will improve the spatial accuracy of the C3PO reconstruction (for example longer exposure times or more powerful lasers). For grid-mode C3PO, a FACS machine capable of multi-way sorting into more tube/wells simultaneously will increase the the number of cells recovered for each tube/well and thus improve the signal-to-noise ratio in any following RNA-Seq. At the stage of library preparation and sequencing, a deeper sampling of the transcriptome could potentially improve the accuracy of the C3PO reconstruction. Furthermore, linear amplification of RNA/cDNA, template switching via random hexamers instead of a template switching oligo and a better correction of PCR duplicates with the addition of UMIs to the sequences before any amplification is performed should all result in more accurate reads counts. Finally, at the stage of reconstruction, knowledge about the tissue configuration can be used to improve the spatial accuracy of C3PO. For example the pre-dissociation light-sheet image can provide useful geometric information about where cells could have originated in 3D space allowing us to position them more accurately. Additionally, algorithmic approaches such as PCA, and other dimensionality-reduction techniques, could help to enforce the assumption that cells in physical proximity share similar gene expression profiles, to attempt to improve spatial reconstruction.

As described in the introduction, a distinct approach commonly used to reconstruct spatial gene expressions is to integrate data from multiple samples - often described as creating an atlas. This requires the samples to be considered substantially identical to each other - for example multiple embryos of the exact same developmental stage. It cannot therefore be used for unique samples like tumours, or rare mutants. A common implementation of this strategy has been to combine deeper sampled single-cell omics data with imaging or spatial omics techniques revealing their positions in space^38^. For example a recent gene expression atlas has been constructed for human foetal limbs by combining Visium with scRNASeq^39^. Many types of information can be integrated in this way such as marker gene expression patterns^5,40–42^ and the organization of cell types in space^42,43^. In a similar fashion such information could be integrated with C3PO spatial information to generate a more accurate hybrid spatial transcriptomic map.

To summarize, to our knowledge we have demonstrated proof-of-principle of the first tool to preserve the original 3D spatial information of an entire tissue after its cells have been dissociated. C3PO can in theory be used in ‘plug-and-play’ fashion with any type of omics downstream (not only the ‘C3PO-Seq’ that we have presented here). Furthermore, we purposely designed the C3PO protocol so that it could be used on any tissue from any species - it does not depend on transgenically-expressed fluorescent proteins. C3PO should prove to be a powerful tool not only for performing omics on wildtype specimens but also mutants such as cancers where marker gene expression typically isn’t available. In cancer biology multiple features could be elucidated using C3PO including the spatial configuration of distinct tumour subtypes of heterogeneous tumours, the distribution of Tumour T-cell infiltration and clonality, and neoantigen distribution.

## Acknowledgments

We thank Xavier Diego and Ben Lehner for helpful discussions. We thank Fumio Nakaki and Antoni Matyjaszkiewicz for reviewing the manuscript. We thank the following core facilities that were used to generate the data in the manuscript: EMBL Mesoscopic Imaging Facility (EMBL Barcelona), UPF/CRG Flow Cytometry Unit, UPF Genomics and EMBL Genecore. Author contributions: Original concept was by J.Swoger. Experimental/conceptual development was performed by J.C, J.Sharpe and J.Swoger. Embryology was performed by J.C and H.C. HCR was performed by A.R. All other experiments were performed by J.C. Analysis was performed by J.C, M.M and J.Sharpe. The manuscript was written by J.C, J.Sharpe and J.Swoger. Funding acquisition by J.Sharpe, J.C and M.M. Supervision by J.Sharpe. Funding: European Research Council Advanced Grant (SIMBIONT, Project no. 670555). Spanish AEI Plan Estatal grant (LIMBNET3D, PID2019-110868GB-I00). Core funding from EMBL and CRG. Competing interests: J.C, J.Sharpe, J.Swoger and A.R have an ongoing international patent application for C3PO with application number PCT/EP2023/087544.

## List of Supplementary Materials

Materials and Methods

Supplementary text, figures and tables

S1) Calibration graphs for bleaching the 3 fluorophores used in this manuscript

S2) Proof-of-principal of pure C3PO with all-or-nothing bleaching

S3) Repeats of pure C3PO reconstructing a mouse limb bud in 3D

S4) 1D spatial transcriptomics of the mouse limb bud

S5) Repeat of 2D spatial transcriptomics of the mouse limb bud

S6) Unbiased analysis of C3PO 2D spatial transcriptomics result

S7) Tissue dimension sizes for mapping fluorescence space to physical space

S8) C3PO spatial precision

Links to Creative Commons licences

Supplementary Movie S1

## References and Notes

1. Tang, F. et al. mRNA-Seq whole-transcriptome analysis of a single cell. Nat. Methods 6, 377–382 (2009).

2. Bandura, D. R. et al. Mass Cytometry: Technique for Real Time Single Cell Multitarget Immunoassay Based on Inductively Coupled Plasma Time-of-Flight Mass Spectrometry. Anal. Chem. 81, 6813–6822 (2009).

3. Stoeckius, M. et al. Simultaneous epitope and transcriptome measurement in single cells. Nat. Methods 14, 865–868 (2017).

4. Rao, A., Barkley, D., França, G. S. & Yanai, I. Exploring tissue architecture using spatial transcriptomics. Nature 596, 211–220 (2021).

5. Achim, K. et al. High-throughput spatial mapping of single-cell RNA-seq data to tissue of origin. Nat. Biotechnol. 33, 503–509 (2015).

6. Richardson, L. et al. EMAGE mouse embryo spatial gene expression database: 2014 update. Nucleic Acids Res. 42, D835–D844 (2014).

7. Lein, E. S. et al. Genome-wide atlas of gene expression in the adult mouse brain. Nature 445, 168–176 (2007).

8. Lubeck, E., Coskun, A. F., Zhiyentayev, T., Ahmad, M. & Cai, L. Single-cell in situ RNA profiling by sequential hybridization. Nat. Methods 11, 360–361 (2014).

9. Shah, S., Lubeck, E., Zhou, W. & Cai, L. In Situ Transcription Profiling of Single Cells Reveals Spatial Organization of Cells in the Mouse Hippocampus. Neuron 92, 342–357 (2016).

10. Eng, C.-H. L. et al. Transcriptome-scale super-resolved imaging in tissues by RNA seqFISH+. Nature 568, 235–239 (2019).

11. Chen, K. H., Boettiger, A. N., Moffitt, J. R., Wang, S. & Zhuang, X. Spatially resolved, highly multiplexed RNA profiling in single cells. Science 348, aaa6090 (2015).

12. Codeluppi, S. et al. Spatial organization of the somatosensory cortex revealed by osmFISH. Nat. Methods 15, 932–935 (2018).

13. Bio-Techne Announces Commercial Release Of RNAscope® HiPlex Assay: A Multiplex In Situ Hybridization Assay For Tissues :: Bio-Techne Corporation (TECH). Bio-Techne Corporation https://investors.bio-techne.com/press-releases/detail/148/bio-techne-announces-commercial-release-of-rnascope (2019).

14. Gyllborg, D. et al. Hybridization-based in situ sequencing (HybISS) for spatially resolved transcriptomics in human and mouse brain tissue. Nucleic Acids Res. 48, e112 (2020).

15. Lee, J. H. et al. Fluorescent in situ sequencing (FISSEQ) of RNA for gene expression profiling in intact cells and tissues. Nat. Protoc. 10, 442–458 (2015).

16. Chen, X., Sun, Y.-C., Church, G. M., Lee, J. H. & Zador, A. M. Efficient in situ barcode sequencing using padlock probe-based BaristaSeq. Nucleic Acids Res. 46, e22 (2018).

17. Wang, X. et al. Three-dimensional intact-tissue sequencing of single-cell transcriptional states. Science 361, eaat5691 (2018).

18. Rodriques, S. G. et al. Slide-seq: A scalable technology for measuring genome-wide expression at high spatial resolution. Science (2019) doi:10.1126/science.aaw1219.

19. Vickovic, S. et al. High-definition spatial transcriptomics for in situ tissue profiling. Nat. Methods 16, 987–990 (2019).

20. Ståhl, P. L. et al. Visualization and analysis of gene expression in tissue sections by spatial transcriptomics. Science 353, 78–82 (2016).

21. Chen, A. et al. Spatiotemporal transcriptomic atlas of mouse organogenesis using DNA nanoball-patterned arrays. Cell 185, 1777-1792.e21 (2022).

22. Cho, C.-S. et al. Microscopic examination of spatial transcriptome using Seq-Scope. Cell 184, 3559-3572.e22 (2021).

23. Wang, X. et al. Three-dimensional intact-tissue sequencing of single-cell transcriptional states. Science 361, eaat5691 (2018).

24. Medaglia, C. et al. Spatial reconstruction of immune niches by combining photoactivatable reporters and scRNA-seq. Science 358, 1622–1626 (2017).

25. Bastide, S. et al. TATTOO-seq delineates spatial and cell type–specific regulatory programs in the developing limb. Sci. Adv. 8, eadd0695 (2022).

26. Kishi, J. Y. et al. Light-Seq: light-directed in situ barcoding of biomolecules in fixed cells and tissues for spatially indexed sequencing. Nat. Methods 19, 1393–1402 (2022).

27. Mai, H. & Lu, D. Tissue clearing and its applications in human tissues: A review. VIEW n/a, 20230046.

28. Optical Sectioning Deep Inside Live Embryos by Selective Plane Illumination Microscopy | Science. https://www.science.org/doi/10.1126/science.1100035.

29. Mayer, J. et al. OPTiSPIM: integrating optical projection tomography in light sheet microscopy extends specimen characterization to nonfluorescent contrasts. Opt. Lett. 39, 1053–1056 (2014).

30. Lewandowski, J. P. et al. Spatiotemporal regulation of GLI target genes in the mammalian limb bud. Dev. Biol. 406, 92–103 (2015).

31. Zhu, J., Patel, R., Trofka, A., Harfe, B. D. & Mackem, S. Sonic hedgehog is not a limb morphogen but acts as a trigger to specify all digits in mice. Dev. Cell 57, 2048-2062.e4 (2022).

32. Probst, S. et al. SHH propagates distal limb bud development by enhancing CYP26B1-mediated retinoic acid clearance via AER-FGF signalling. Development 138, 1913–1923 (2011).

33. Matsubara, Y. et al. Inactivation of Sonic Hedgehog Signaling and Polydactyly in Limbs of Hereditary Multiple Malformation, a Novel Type of Talpid Mutant. Front. Cell Dev. Biol. 4, (2016).

34. Yokoyama, S. et al. A Systems Approach Reveals that the Myogenesis Genome Network Is Regulated by the Transcriptional Repressor RP58. Dev. Cell 17, 836–848 (2009).

35. Montero, J. A. et al. Sox9 Expression in Amniotes: Species-Specific Differences in the Formation of Digits. Front. Cell Dev. Biol. 5, (2017).

36. McGlinn, E. et al. Expression of the NET family member Zfp503 is regulated by hedgehog and BMP signaling in the limb. Dev. Dyn. 237, 1172–1182 (2008).

37. Markman, S. et al. A single-cell census of mouse limb development identifies complex spatiotemporal dynamics of skeleton formation. Dev. Cell 58, 565-581.e4 (2023).

38. Longo, S. K., Guo, M. G., Ji, A. L. & Khavari, P. A. Integrating single-cell and spatial transcriptomics to elucidate intercellular tissue dynamics. Nat. Rev. Genet. 22, 627–644 (2021).

39. Zhang, B. et al. A human embryonic limb cell atlas resolved in space and time. Nature 1–11 (2023) doi:10.1038/s41586-023-06806-x.

40. Puram, S. V. et al. Single-Cell Transcriptomic Analysis of Primary and Metastatic Tumor Ecosystems in Head and Neck Cancer. Cell 171, 1611-1624.e24 (2017).

41. Moor, A. E. et al. Spatial Reconstruction of Single Enterocytes Uncovers Broad Zonation along the Intestinal Villus Axis. Cell 175, 1156-1167.e15 (2018).

42. Moriel, N. et al. NovoSpaRc: flexible spatial reconstruction of single-cell gene expression with optimal transport. Nat. Protoc. 16, 4177–4200 (2021).

43. Saunders, A. et al. Molecular Diversity and Specializations among the Cells of the Adult Mouse Brain. Cell 174, 1015-1030.e16 (2018).

44. Musy, M. et al. A quantitative method for staging mouse embryos based on limb morphometry. Dev. Camb. Engl. 145, dev154856 (2018).

45. Babraham Bioinformatics - FastQC A Quality Control tool for High Throughput Sequence Data. https://www.bioinformatics.babraham.ac.uk/projects/fastqc/.

46. Martin, M. Cutadapt removes adapter sequences from high-throughput sequencing reads. EMBnet.journal 17, 10–12 (2011).

47. Langmead, B. & Salzberg, S. L. Fast gapped-read alignment with Bowtie 2. Nat. Methods 9, 357–359 (2012).

48. Danecek, P. et al. Twelve years of SAMtools and BCFtools. GigaScience 10, giab008 (2021).

49. Liao, Y., Smyth, G. K. & Shi, W. The Subread aligner: fast, accurate and scalable read mapping by seed-and-vote. Nucleic Acids Res. 41, e108 (2013).

50. Musy, M. et al. marcomusy/vedo: 2023.5.0. Zenodo 10.5281/zenodo.4587871 (2023).

51. Zhou, W. et al. Misexpression of Pknox2 in Mouse Limb Bud Mesenchyme Perturbs Zeugopod Development and Deltoid Crest Formation. PLOS ONE 8, e64237 (2013).

52. Drake, K. D., Lemoine, C., Aquino, G. S., Vaeth, A. M. & Kanadia, R. N. Minor spliceosome disruption causes limb growth defects without altering patterning. 2020.03.16.994384 Preprint at 10.1101/2020.03.16.994384 (2020).

53. Tissières, V. et al. Gene Regulatory and Expression Differences between Mouse and Pig Limb Buds Provide Insights into the Evolutionary Emergence of Artiodactyl Traits. Cell Rep. 31, 107490 (2020).

54. Pennimpede, T., Cameron, D. A., MacLean, G. A. & Petkovich, M. Analysis of Cyp26b1/Rarg compound-null mice reveals two genetically separable effects of retinoic acid on limb outgrowth. Dev. Biol. 339, 179–186 (2010).

55. Beverdam, A. & Meijlink, F. Expression patterns of group-I aristaless-related genes during craniofacial and limb development. Mech. Dev. 107, 163–167 (2001).

56. Kuhlbrodt, K. et al. Cooperative Function of POU Proteins and SOX Proteins in Glial Cells *. J. Biol. Chem. 273, 16050–16057 (1998).

57. Goldstein, J. M. et al. Variation in zygotic CRISPR/Cas9 gene editing outcomes generates novel reporter and deletion alleles at the Gdf11 locus. Sci. Rep. 9, 18613 (2019).

58. Gurniak, C. B., Perlas, E. & Witke, W. The actin depolymerizing factor n-cofilin is essential for neural tube morphogenesis and neural crest cell migration. Dev. Biol. 278, 231–241 (2005).

59. De Arcangelis, A., Georges-Labouesse, E. & Adams, J. C. Expression of fascin-1, the gene encoding the actin-bundling protein fascin-1, during mouse embryogenesis. Gene Expr. Patterns 4, 637–643 (2004).

60. van Bueren, K. L. et al. Murine embryonic expression of the gene for the UV-responsive protein p15PAF. Gene Expr. Patterns 7, 47–50 (2007).

61. Flenniken, A. M., Gale, N. W., Yancopoulos, G. D. & Wilkinson, D. G. Distinct and Overlapping Expression Patterns of Ligands for Eph-Related Receptor Tyrosine Kinases during Mouse Embryogenesis. Dev. Biol. 179, 382–401 (1996).

62. Pellegrini, M., Pantano, S., Fumi, M. P., Lucchini, F. & Forabosco, A. Agenesis of the Scapula in Emx2 Homozygous Mutants. Dev. Biol. 232, 149–156 (2001).

63. Wehrle-Haller, B., Morrison-Graham, K. & Weston, J. A. Ectopic c-kit Expression Affects the Fate of Melanocyte Precursors inPatchMutant Embryos. Dev. Biol. 177, 463–474 (1996).

64. Keeney, D. S. & Waterman, M. R. Two novel sites of expression of NADPH cytochrome P450 reductase during murine embryogenesis: Limb mesenchyme and developing olfactory neuroepithelia. Dev. Dyn. 216, 511–517 (1999).

65. Itou, J. et al. HMGB factors are required for posterior digit development through integrating signaling pathway activities. Dev. Dyn. 240, 1151–1162 (2011).

