## Supplementary Materials for "*C*ell *3*D *P*ositioning by *O*ptical encoding *(C3PO*) and its application to spatial transcriptomics"

### Materials and Methods

#### Labelling tissue

Tissue was fixed with 4% PFA shaking at room temperature for 1 hour or overnight at 4°C on a shaker. Tissue was then permeabilized and endogenous RNAases deactivated by treatment with 0.025% CHAPS, 0.5% Triton with 0.143Molar Beta-mercaptoethanol in PBS overnight at 4°C on a shaker. The sample was quickly washed with PBS and transferred to a PCR tube. Samples were then stained with 10-50μM fluorophore conjugated oligonucleotide with 10% DMSO, 0.01% Triton in PBS overnight in the dark at 4°C on a shaker. The following day the samples were placed in a thermocycler and oligonucleotides bound using 6 cycles of 95°C for 15 seconds and 25°C for 1 minute followed by cooling to 15°C. Samples were then washed in PBS twice quickly and once for approximately 1-2hours in the dark at room temperature on a shaker before being post fixed in 4% PFA for 2 hours in the dark at room temperature on a shaker before finally being moved to PBS.

#### Confocal imaging to determine nuclear location of signal

Limb buds were stained in the same way as previously described but with only the oligonucleotide covalently attached to 6FAM and Alexa647 in addition to 1 in 5000 DAPI. Limb buds were imaged on a mattek dish using an inverted SP5 Microscope (Leica). DAPI, 6FAM and Alexa647 fluorophores were excited with 405nm, 488nm and 633nm laser lines respectively. Fluorescence was detected with PMT ranges spanning the emission spectrum of the respective fluorophores.

#### Mounting, dehydrating and clearing samples

Samples were quickly washed in milliQ water before being mounted in 1% low melting point agarose dissolved in Milliq water and filtered through a 0.45μm filter. After mounting, samples were dehydrated by washing in 100% methanol once quickly and twice for >4hours in the dark in glass vials at 4°C. Samples were then cleared by washing in 1:2 benzyl alcohol:benzyl benzoate (BABB), once quickly and twice >4hours in the dark in glass vials at 4°C.

#### Multiplexed in situ Hybridization Chain Reaction (HCR$^{TM}$)

Fluorescent In Situ HCR$^{TM}$ was performed following Molecular Instruments' protocol for mouse embryos (available at www.molecularinstruments.com). In brief, sox9, hoxa11 and hoxa13 HCR$^{TM}$ probe sets were designed and purchased from Molecular Instruments and let hybridized to RNA targets by incubating overnight at 37ºC. After removing excess of probe by several washes, mouse limbs were pre-amplified with amplification buffer and incubated overnight at room temperature with h1 and h2 hairpins that create an amplification polymer coupled with the corresponding fluorophore. In this specific case, hoxa11 was designed for use with HCR amplifier B3 coupled with 488 fluorophore, hoxa13 for use with HCR amplifier B2 coupled with 546 fluorophore and sox9 for use with HCR amplifier B1 coupled with 647 fluorophore. E10.5 mouse limb was embedded in agarose and all three genes were imaged in the MuVI SPIM light-sheet microscope from Luxendo, Bruker.

#### Imaging and photo-bleaching

Samples were imaged and bleached on an custom-built OPTiSPIM light sheet mesoscope[29]. Briefly, a 50mW 405nm, a 50mW 488nm, and a 50mW 639nm lasers were used to image and bleach the Alexa405, 6FAM and Alexa647 fluorophores respectively. A 5mW 543nm HeNe laser was used to image the staining uniformity control TAMRA fluorophore. 447BP60, 525BP50, 585BP60, and 700BP75 band pass filters were used for detecting the Alexa405, 6FAM, TAMRA and Alexa647 fluorophores respectively. A 2.5× N Plan air objective (Leica, NPLAN, NA=0.07) was used to illuminate the samples. A 5× N Plan Epi air objective (Leica, NPLAN EPI, NA=0.12, WD=14 mm) and a 12 bit cooled Hamamatsu ORCA-ER C4742-80 CCD camera were used for detection.

A calibration sample was used to calibrate the bleaching, by placing the sample at different positions relative to the light sheet for each laser line and monitoring the reduction in fluorescent signal with time of exposure at 100% laser power (see SM Fig. S1). Bleaching exposure times for different positions of the sample of interest for C3PO analysis are determined through reading off of these calibration graphs (one for each fluorophore/laser line).

**Processing of photo-bleached samples**

After bleaching, the sample received one quick wash and two longer washes (>4hours) in methanol at 4°C. The sample was then removed from the LMP agarose and rehydrated with one quick wash and two longer washes (>4 hours) in PBS at 4°C. For the experiments where the limb ectoderm and mesenchyme are analyzed separately the ectoderm was removed from the limb with tweezers after incubating the limb bud in 0.5% Trypsin-EDTA 10x (without phenol red - Gibco 15400-054) for 15 minutes at room temperature. For RNA-based experiments the tissue was digested with Tryple Select (with $1/1x10^6$ Triton) and collagenase type III at 37°C 1200 RPM for 2 hours. For any other experiment the tissue was digested with 0.22µm filtered 10mg/ml Collagenase/dispase in PBS (with $1/1x10^6$ Triton) at 37°C 600 RPM for 2 hours. Tissue samples were periodically mechanically agitated with a gilson pipette. Following tissue dissociation, the sample was centrifuged at 600g for 5 minutes at room temperature. Cell pellets were resuspended in 0.5-1ml PBS and strained through a 40µm strainer and processed on a FACS analyzer (BD Fortessa) or sorter (BD FACSAria II).

**FACS**

For experiments where cells were sorted and RNA isolated a FACSAria cell sorter was used. For experiments where cells were simply analysed for their fluorescent levels the Fortessa FACS analyzer was used. To detect Alexa405 fluorophores a 405nm laser with 450BP50 bandpass filter was used on the Fortessa and a 407nm laser with 450BP40 bandpass filter was used on the FACSAria. To detect 6FAM fluorophores a 488nm laser with 530BP28 bandpass filter and 505LP longpass filter was used on the Fortessa and a 488nm laser with 525BP50 bandpass filter and 505LP longpass filter was used on the FACSAria. To detect TAMRA fluorophores a 561nm laser with 586BP15 or 610BP20 bandpass filter was used on the Fortessa and a 561nm laser with 585BP15 or 610BP20 bandpass filter was used on the FACSAria. To detect Alexa647 fluorophores a 633nm laser with 670BP14 bandpass filter was used on the Fortessa and a 633nm laser with 670BP14 bandpass filter was used on the FACSAria. On both machines, a FACS gate for Side scatter area and Forward scatter was used to remove debris (P1). Singlets were then selected with a Forward scatter height versus area gate (P2). A PE-A (with similar fluorescence profile to TAMRA) versus Forward scatter gates was used to select the fluorescent cells and remove any remaining debris (P3).

PACB (corresponding to Alexa405), FITC-A (corresponding to 6FAM) and APC-A (corresponding to Alexa647) fluorescent channel levels were normalized to the PE-A (corresponding to TAMRA) fluorescent channel level to generate the data shown in Fig. 3.

**Grid-mode transcriptomics**

For 2D grid mode C3PO the tissue was bleached using gradients for 2 axes, using 639nm bleaching for the proximal-distal axis and 405nm bleaching for the anterior-posterior axis of a forelimb before post-bleach imaging performed in all channels. The tissue was then processed as described above but with the following modification for the cell sorting downstream of the TAMRA+ gate: A 2D grid of gates (14 or 15) was set up in a Alexa647/TAMRA ratio versus Alexa405/TAMRA ratio plot. Cells were then sorted into individual Eppendorf tubes that correspond to each specific gate. For the 1D proximal-distal grid mode C3PO Cells were sorted based on the relative level of APC-A (corresponding to Alexa647) to PE-A (corresponding to TAMRA) into 4 separate Eppendorf tubes corresponding to proximal, mid-proximal, mid-distal and distal cells (see SM section S4).

To determine which side of the limb was anterior and which was posterior for the 2D grid mode C3PO a double validation approach was utilised using 1) the EMOSS[44] limb staging system which allows the determination of whether the dorsal or ventral side of the limb is being viewed. 2) 3D projections of the limb using the TAMRA unbleached control channel were made in ImageJ. The 3D projection allowed the observation of the curvature of the limb relative to the flank (it typically curves inwards towards the centre of the body) which allowed the determination of which side of the limb was dorsal and which was ventral. Using the knowledge of which side of the limb is dorsal/ventral and the knowledge of whether a left or right limb was being processed it is possible to determine which side of the limb is anterior and which is posterior. The two methods had to agree for us to continue with the experiment.

**RNA Extraction**

Cells were sorted into 45µl RLT buffer (Qiagen RNAEasy kit) and 5µl ProteinaseK (Qiagen Blood and tissue kit). Samples were then heated at 56°C for 1 hour shaking at 1200 RPM. 125µl of RNA clean beads (Beckman Coulter) were added to the sample, vigorously mixed and let stand at room temperature for 5 minutes. Samples were then placed on a magnet for 5 minutes and washed twice with 70% Ethanol for 30 seconds each. Pellets were air dried for 10 minutes on the magnet before being removed. Beads were resuspended in 15µl of nuclease free water and vigorously mixed. Samples were stood at room temperature for 5 minutes and then placed on a magnet. After 5 minutes the RNA was removed from the tubes.

**Library preparation**

Libraries were prepared using the NEBNext single cell/Low input RNA kit. However 1.0x SPRISelect beads (Beckman Coulter) were used instead of 0.6x beads and 21 rounds of amplification performed on cDNA libraries. Libraries were run on a Nextseq 2000 on High output mode using 100bp single or paired end reads.

**Bioinformatics**

Bioinformatics was carried out on a HPC cluster using the SLURM scheduling system. Each fastq sequence file was processed by a different CPU as a separate job. Fastqc[45] was then used to confirm read quality. Modules were then loaded for cutadapt, bowtie2, SAMtools and

Subread. Two rounds of cutadapt[46] were then utilised to first trim the NEB multiplex sequencing oligonucleotide sequence and then trim the NEBNext template switching oligonucleotide and first strand synthesis oligonucleotide sequences using the following parameters:

cutadapt -a AGATCGGAAGAGCACACGTCTGAACTCCAGTCA

cutadapt -a AAGCAGTGGTATCAACGCAGAGTACTTTTTTTTTTTTTTTTTTTTTTTTT -g GCTAATCATTGCAAGCAGTGGTATCAACGCAGAGTACATGGG

Bowtie2[47] (in local mode and single end or paired end mode depending on the dataset) was used to map the resulting trimmed sequences to the mouse mm10 reference genome. Samtools[48] was used to convert data to bam format, remove the PCR duplicates and sort/index the data using the fixmate, sort, markdup and index functions.

In order to count the incidence of the mRNAs Subread's[49] featureCounts function was used. First the gtf annotation file for mouse genome mm10 was downloaded from:

http://hgdownload.soe.ucsc.edu/goldenPath/mm10/bigZips/genes/mm10.refGene.gtf.gz

Then featureCounts was run with the following parameters:

-p -Q 30 -a mm10.refGene.gtf -t 3UTR -g gene_id -o counts.txt sample.bam,

in order to process only paired end reads that map to the 3'UTR (using the -t 3UTR parameter flag) with phred score above 30 (using the -Q 30 parameter). Read counts for each gene are then normalized to the total number of reads for that particular sample.

**Tissue reconstruction, rendering and gene expression mapping:**
In general tissue was reconstructed and rendered using the vedo toolkit[50]. FACS measured fluorescent level data for the channels corresponding to the 3 Cartesian coordinate axes were normalized to the internal labelling control channel. The normalized data was then read as a point cloud in python/vedo.

To generate the pure C3PO (with no transcriptomics) point cloud renders shown in Fig. 3 and Fig. S3 vedo's remove_outliers function was used with radius=0.1 and neighbours=5.

To reconstruct the ectoderm surface shown in Fig. 3d we first scaled the point cloud data to fit a unit cube, then vedo's "remove_outliers" function was used with parameters radius=0.05 and neighbours=5. The resulting point cloud was then sub-sampled (fraction=0.01) and smoothed (with smoothing factor=2). A polygonal mesh was finally generated using the method "reconstruct_surface" (with parameters dims=50, radius=0.13) and the largest region extracted using method "extract_largest_region".

To generate the 2D tissue reconstructions shown in Fig. 4, Fig. S5 and Fig. S6 cell positions for each of the sorted gates were extracted from the FACS data using FCS Express. Gene expression values were plotted for individual cells according to their corresponding FACS gate with the aim of utilizing the full dynamic range of vedo's blue colour map.

### Supplementary text, figures and tables

#### S1) Calibration graphs for bleaching the 3 fluorophores used in this manuscript

A mouse limb bud at approximately E10.5 of age was used to calibrate the bleaching, by placing each laser line lightsheet in a fixed position and monitoring the reduction in intensity of fluorescent signal with time of exposure. The intensity versus time of exposure plots can be seen in Fig. S1. The reduction in intensity does not follow an exponential decay type distribution, instead bleaching saturates over time. For this reason bleaching times for generating gradients of bleaching were manually read off of calibration graphs instead of calculated.

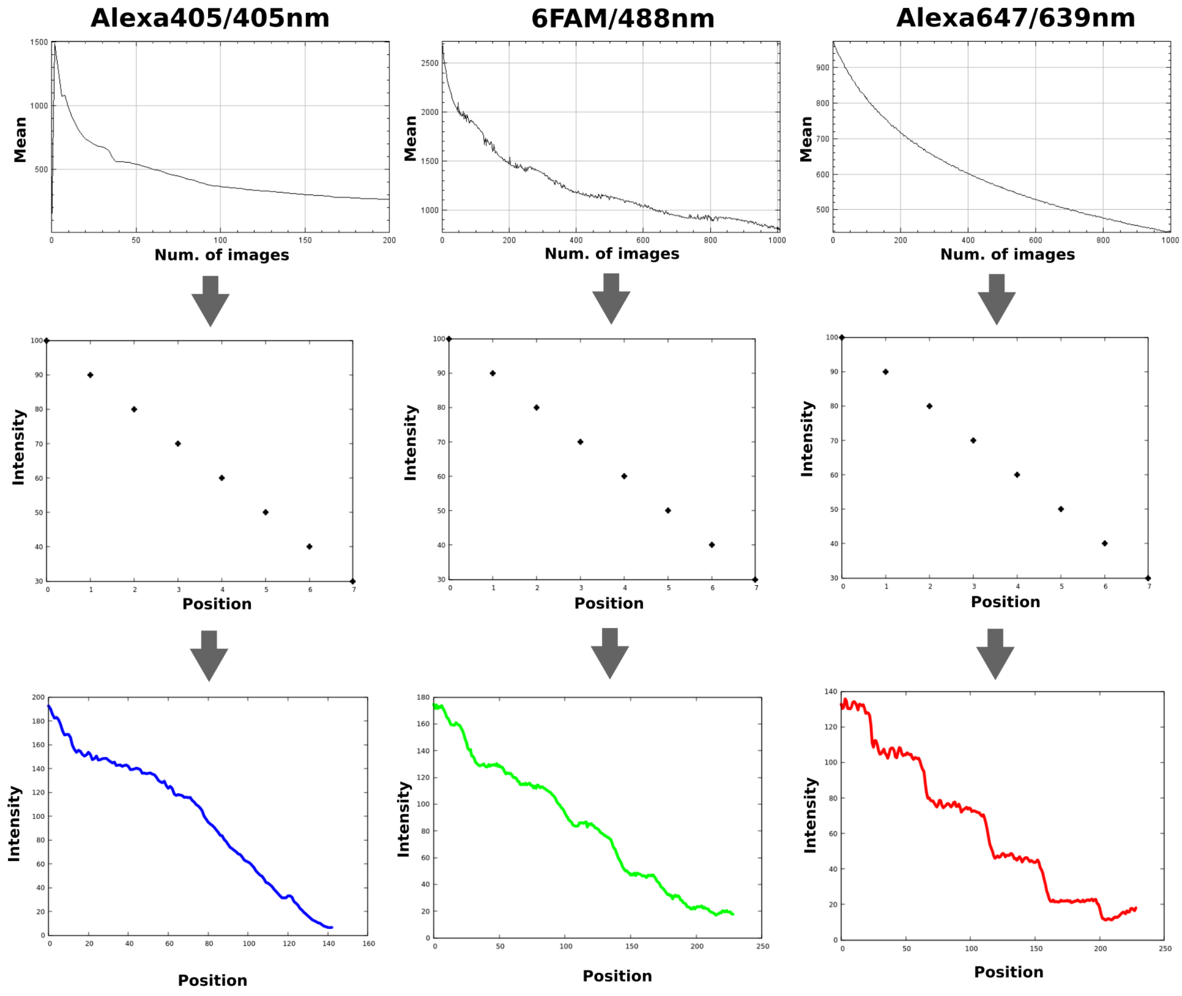


**Figure S1: Calibration of bleaching.** The light sheet for each laser line was parked in a single position in the tissue and time lapse imaged. Camera settings were adjusted so as not to overexpose and saturate the pixel value of any region of the tissue. The average intensity of the image is plotted over time. For Alexa405 a 0.05 second exposure of the 405nm laser at 100% laser power was used at 10 milliwatts. For 6FAM a 0.8 second exposure of the 488nm laser at 100% laser power was used at 50 milliwatts. For Alexa647 a 0.6 second exposure of the 639nm laser at 100% laser power was used at 50 milliwatts.

<u>**S2) Proof of principal of pure C3PO with All-or-nothing bleaching**</u>

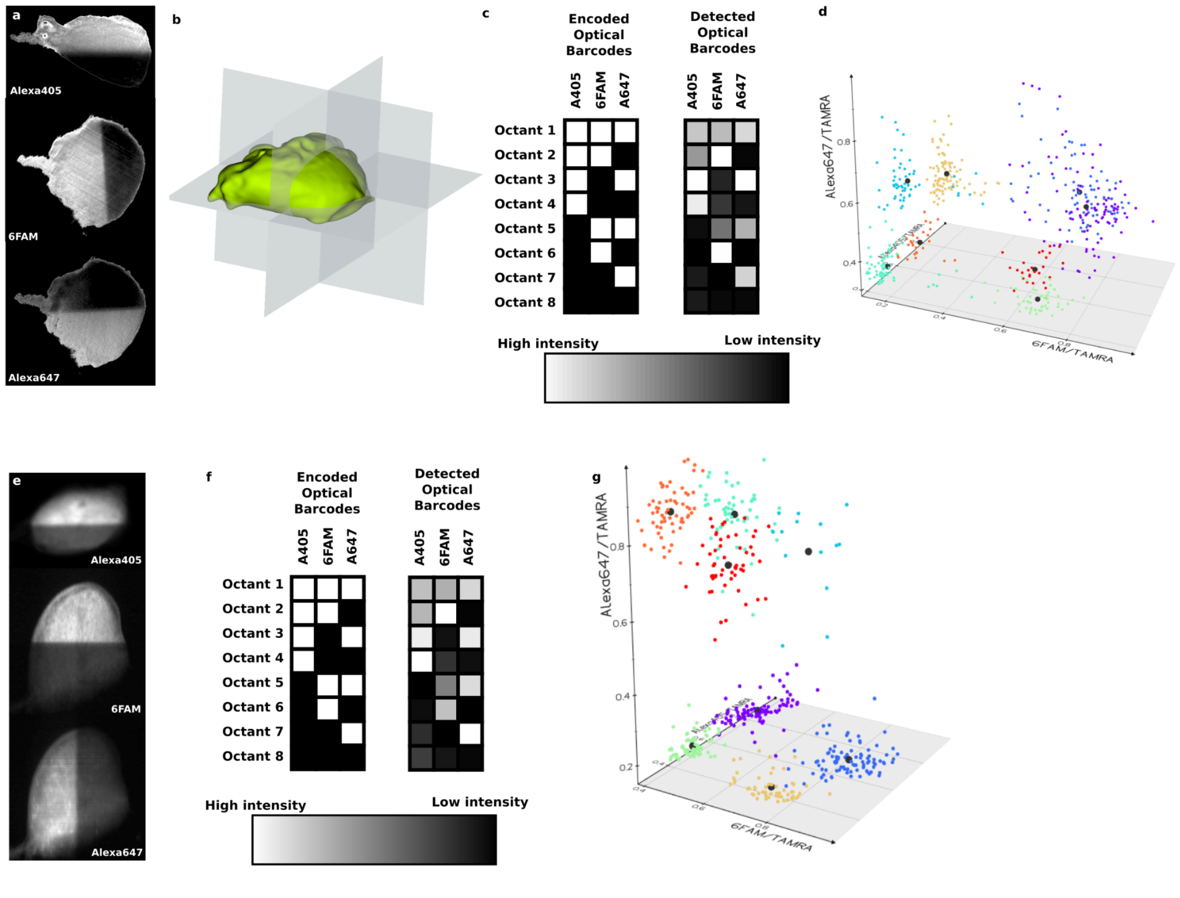


**Figure S2: Proof-of-principle of pure C3PO.** (a) All-or-nothing bleaching of a developing mouse limb bud in the 3 orthogonal axes. Sum-projection images show the respective bleached channel normalized to the labelling uniformity control channel after subtraction of background signal (measured by the mode of the image stack). (b) Such bleaching of the limb bud encodes distinct optical addresses into each of 8 octant regions. Here, we term the optical addresses ‘optical barcodes’ since they are constructed of digital all-or-nothing signals. (c and d) The limb bud was dissociated to single cells and the fluorescent signals measured via FACS. The scatter plot shows 500 randomly chosen cells where each axis corresponds to the measured signal in one of the bleached channels normalized to the labelling uniformity control channel. The cells were hierarchically clustered (see methods) according to the normalized levels of the 3 channels with 8 clusters selected. The colour of the points in the scatter graph indicates the cluster they were assigned to and the large point represents the centroid of each cluster. The normalized fluorescent levels of the 8 centroids were used to determine the detected optical barcodes shown in (c). The encoded and detected optical barcodes were aligned to be as similar as possible demonstrating that the correct pattern of optical barcodes is detected. (e-g) A repeat of the previous experiment again elucidating 8 populations with distinct optical barcodes as encoded demonstrating the consistency of the technique.

### S3) Repeats of pure C3PO reconstructing a mouse limb bud in 3D

In Fig. S3 we show the scatter plots for the whole Ectoderm (blue) and Mesenchyme (red) measured via FACS for limbs #2 (left) and #3 (right) for which an ectodermal concavity is shown in Fig. 3.

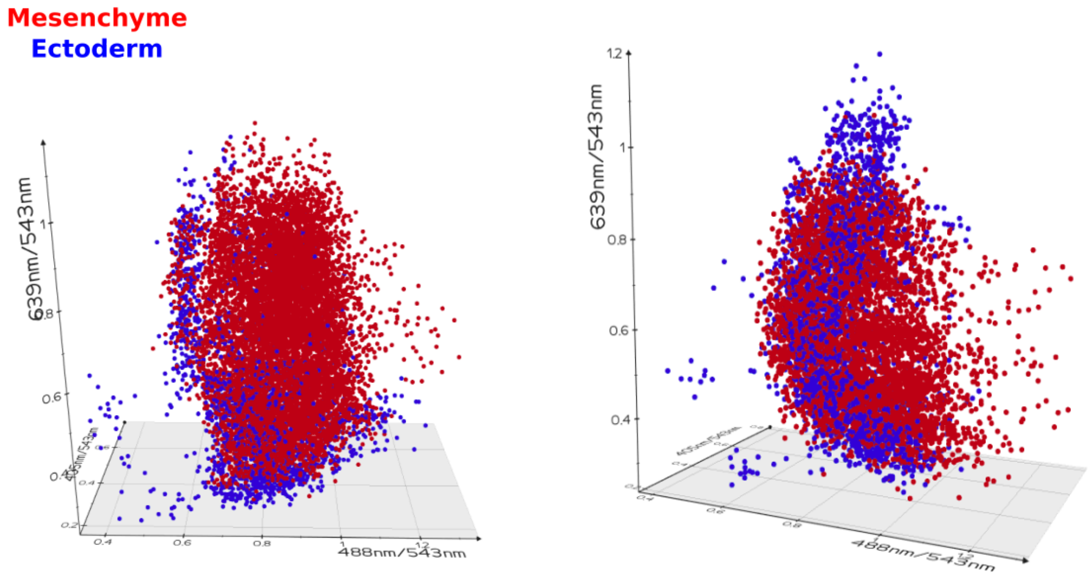


**Figure S3: 2 repeats of the experiment to reconstruct the mouse limb bud tissue configuration.** Ectoderm is shown in blue and mesenchyme is shown in red. A point cloud render of the normalized fluorescence data is shown. In both cases multiple key features of the limb tissue configuration can be reconstructed including 1) The shape of the point clouds resemble a limb shape in the correct orientation relative to the bleaching, 2) The ectoderm is surrounding the mesenchyme, 3) The highest density of ectoderm is around the periphery as expected as this is where the apical ectodermal ridge is located (an ectodermal thickening that acts as a signalling centre during limb development) and 4) The flank regions have a low density of ectoderm. 5) A concavity is observed in the ectoderm (see Fig. 3)

### S4) 1D transcriptomics C3PO

We performed 1-dimensional grid-mode C3PO spatial transcriptomics along the proximal-distance axis of the mouse limb bud (Fig. S4a). We bleached the Alexa647 channel along this axis into 4 separate domains proximal, mid-proximal, mid-distal and distal (Fig. S4b). Tissue was dissociated and FACS sorted into separate tubes depending on the Alexa647 signal relative to the labelling control channel (TAMRA) signal. RNA was extracted from these cells and standard RNASeq library prep performed (see methods). After deep sequencing these libraries, the reads were trimmed, mapped, PCR duplicates removed and reads mapping to the 3' UTR of known genes of the mouse genome counted (see methods). Known control genes such as GAPDH, ActinB and Tubulin-alpha are found to have uniform expression along the PD axis of the limb bud. We then explored the spatial distribution of key genes in proximal-distal patterning with known spatial distributions in the limb bud normalizing to the expression levels of these control genes and compared them to the known gene expression patterns as measured by *in situ* hybridization (Fig. S4c)

The Meis genes including Meis 1 are known to have a high expression in the proximal region of the limb bud and low everywhere else. We find all 3 Meis genes indeed have this profile as shown by the intensity profile of Meis 1 in the 1D C3PO data. FGF signalling read-out genes such as Fgf8 and Dusp6 are known to have high expression in the distal part of the limb bud. Again we find that those genes are expressed in exactly the same pattern in the 1D C3PO data. Certain genes are also known to have a higher expression in the mid region of the proximal-distal axis such as Osr2. We find that those genes also have a peak of expression in the mid regions of the tissue in the 1D C3PO data. Several other key genes important for limb development are also shown. Overall, for the vast majority of genes for which we checked the expression pattern, the 1D C3PO reconstructed pattern was correct.

*Adaptations to protocol described in main manuscript:*
Note that for this experiment there was a slight adaptation in the thermocycler parameters for oligonuclotide binding: 1 cycle of 70C for 30 seconds followed by 25C for 1 minute and cooling to 15C was used. Bioinformatics was performed in the same way as described in the main manuscript but with the following change. After the Fastqc step, the MidLow dataset required quality trimming for which we used `Cutadapt -u -65` to remove low phred score parts of the reads.

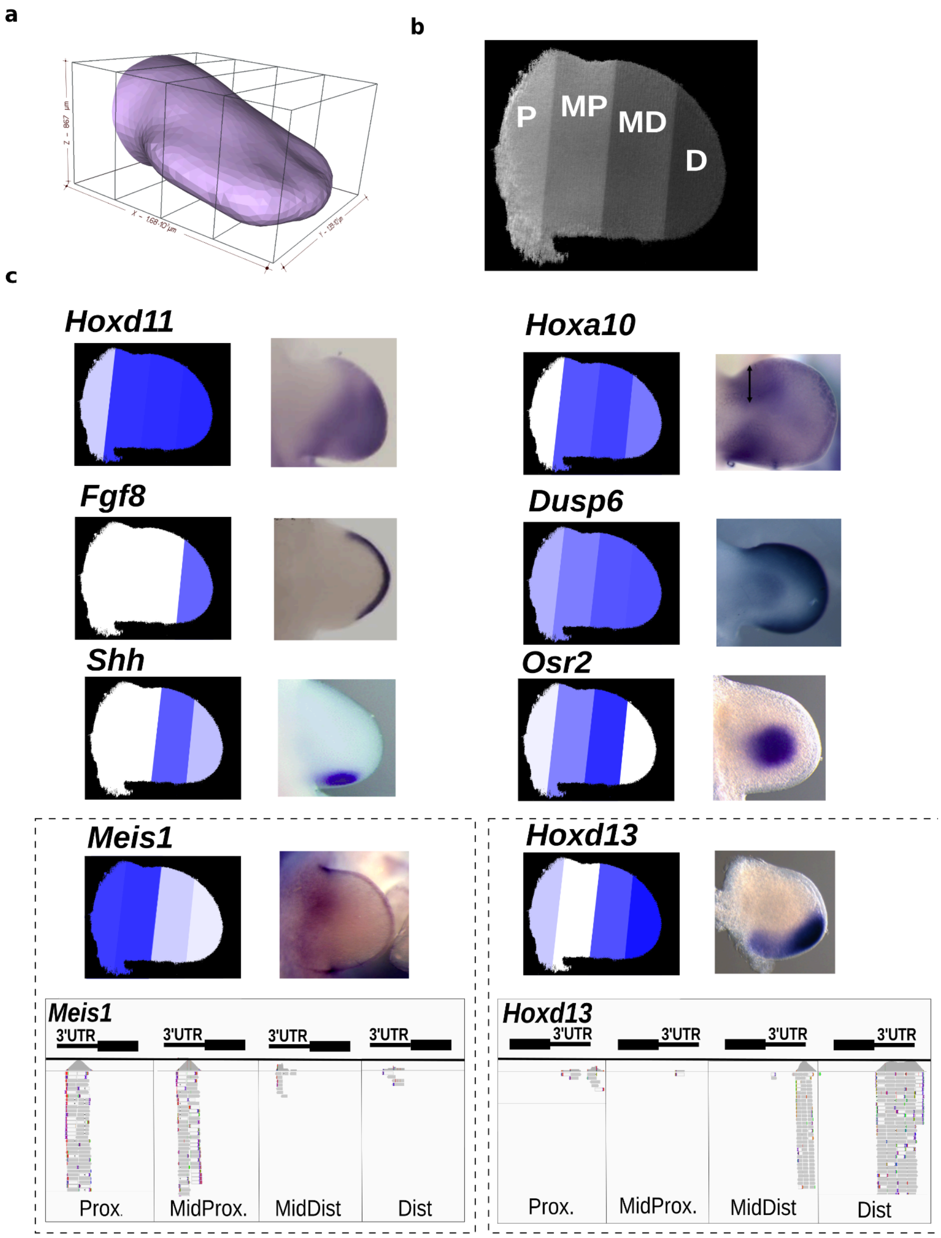


**Figure S4: 1D C3PO spatial transcriptomics.** (a) A schematic of 1D C3PO in grid-mode. (b) Fluorescent signal of Alexa647 fluorophore over the limb bud after bleaching with a 639nm laser is shown. The 4 regions that were sorted in to separate eppendorf tubes (P=proximal, MP=Mid proximal, MD=Mid distal and D=Distal) are shown. (c) On the left is shown the C3PO reconstruction of the gene expression pattern (blue is high expression and white is low). On the right is shown the known gene expression pattern as measured by *in situ* hybridisation. Dashed insets show 2 specific examples of the reads mapped to the mouse genome for the different regions as viewed by Interactive Genomics Viewer (IGV). Whole mount *in situ* hybridisation (WMISH) patterns for *Osr2, Hoxd13,* reprinted from[30] with permission from Elsevier, *Hoxd11 and Fgf8* reprinted from[31] under a Creative Commons CC-BY-NC-ND licence, *Shh* reprinted from[33] under a Creative Commons CC-BY licence, *Dusp6* generated by the Sharpe laboratory, *Meis1* reprinted from the Embrys database[34] with permission from Hiroshi Asahara, *Hoxa10* reprinted from[51] under a Creative Commons CC-BY licence.

### S5) Repeat of 2D spatial transcriptomics of the mouse limb bud

To confirm the results of the 2D spatial transcriptomics shown in the main manuscript we repeated the experiment with an additional limb (limb #5, Fig. S5). Furthermore, we also utilized unbiased scoring of reconstruction of the top expressed 100 genes with an image and a discernible pattern in the MGI or Embrys gene expression databases as shown in Fig. S6.

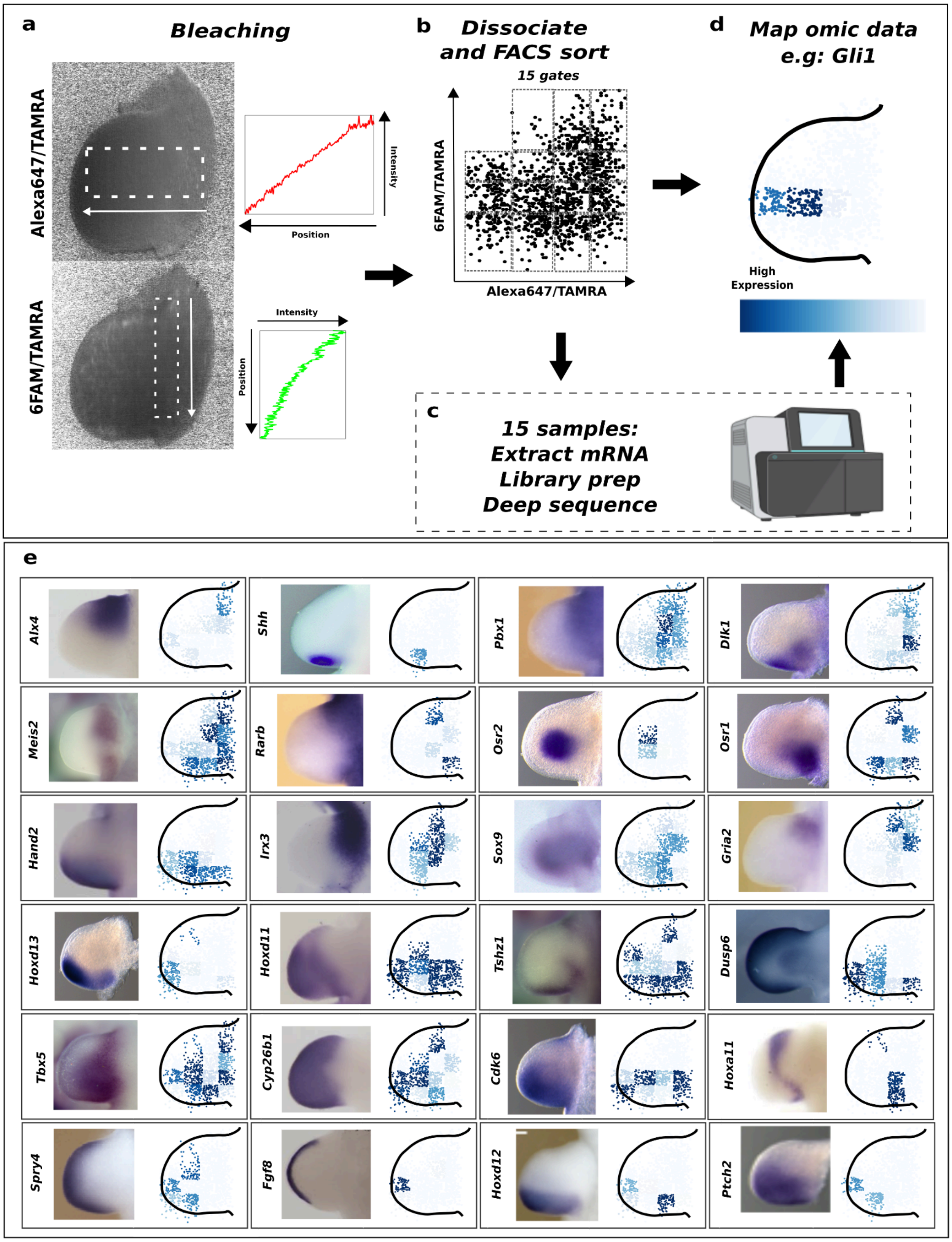

**Figure S5: A repeat of spatial transcriptomics with C3PO in 2D grid-mode** (a) Monotonic gradient bleaching of the 2 principle axes of the developing mouse limb bud: proximal-distal (top) and anterior-posterior (bottom). Sum-projection images show the respective bleached channel normalized to the labelling uniformity control channel. Plots of the signal intensities over spaces corresponding to the dashed boxes for the two channels are shown next to the images. (b) After bleaching the tissue was dissociated and the individual cells were run through a FACS machine. Cells were ratio sorted into 15 different tubes according to their bleached channel signals normalized to the labelling uniformity control channel. The 15 regions were defined so that they encompassed the entire tissue. (c) RNAseq was carried out on the 15 individual samples. (d) That expression data could then be mapped back onto the tissue reconstruction. (e) Examples of C3PO with spatial transcriptomics of 24 key genes involved in mouse limb bud development. The C3PO data is shown next to the known gene expression profile as measured by *in situ* hybridization. Expression level is indicated by the depth of blue shading. Whole mount *in situ* hybridization (WMISH) patterns for *Osr1, Osr2, Dlk1, Hoxd13, Ptch2, Cdk6* reprinted from[30] with permission from Elsevier, *Hand2, Alx4, Irx3, Hoxd11, Cyp26b1, Fgf8* reprinted from[31] under a Creative Commons CC-BY-NC-ND licence, *Rarb, Spry4, Gria2, Pbx1* reprinted from[32] with a licence from Copyright Clearance Center on behalf of The Company of Biologists Ltd., *Shh* reprinted from[33] under a Creative Commons licence, *Sox9* reprinted from[35] under a Creative Commons CC-BY licence, *Dusp6* generated by the Sharpe laboratory, *Tbx5, Tshz1, Meis2* reprinted from the Embrys database[34] with permission from Hiroshi Asahara, *Hoxa11* reprinted from[52] under a Creative Commons CC-BY-ND licence, *Hoxd12* reprinted from[53] under a Creative Commons CC-BY-NC-ND licence.

### S6) Unbiased analysis of C3PO 2D spatial transcriptomics result

To assess C3PO in an unbiased fashion, we combined the results from limb #4 and limb #5. The 100 most expressed genes from each limb bud (from the RNA-Seq results) were added to the 24 genes initially analysed (Fig. 4), resulting in a total of 138 genes (due to overlaps between the lists). We then searched for previously-published expression patterns for all these genes, in two public databases:

1) MGI GXD database (Jackson Laboratory)
    - https://www.informatics.jax.org/gxd
    - Anatomical Structure = “limb bud TS14-19”

2) the embrys database (Tokyo Medical and Dental University).
    - https://www.embrys.jp/embrys/2DView/pictures

87 of these genes had a discernible gene expression pattern, of which 73 (=84%) showed a good or reasonable match with the prediction by C3PO (Fig. S6). Many genes have a distal and/or posterior pattern, while a smaller number show anterior and/or proximal patterns. It is also interesting to distinguish patterns with fairly uniform distributions, from those with complex or very internal patterns. Accuracy can be determined by how similar the C3PO reconstructions match the known gene expression pattern whilst precision can be determined by the similarity of the gene expression pattern reconstructed for the two limbs.

Differences between C3PO and WMISH results may be explained by a few issues:

1) For a few of the genes, the most similar aged image in the databases were slightly older or younger than limb #4 or #5, which could display a slightly different pattern (for example asterisks in Fig. S6 indicate 2 cases that are older).
2) A single cell mis-located into the wrong grid block may occasionally have an exaggerated impact on the whole pattern, due to normalisation effects. This could be improved in the future by suggestions which are explained in the discussion (eg. employing a higher resolution spatial grid, more linear amplification of RNA/cDNAs and addition of UMIs before amplification).
3) It is possible that some WMISH patterns in the databases were done with a probe representing a different isoform of the transcript.

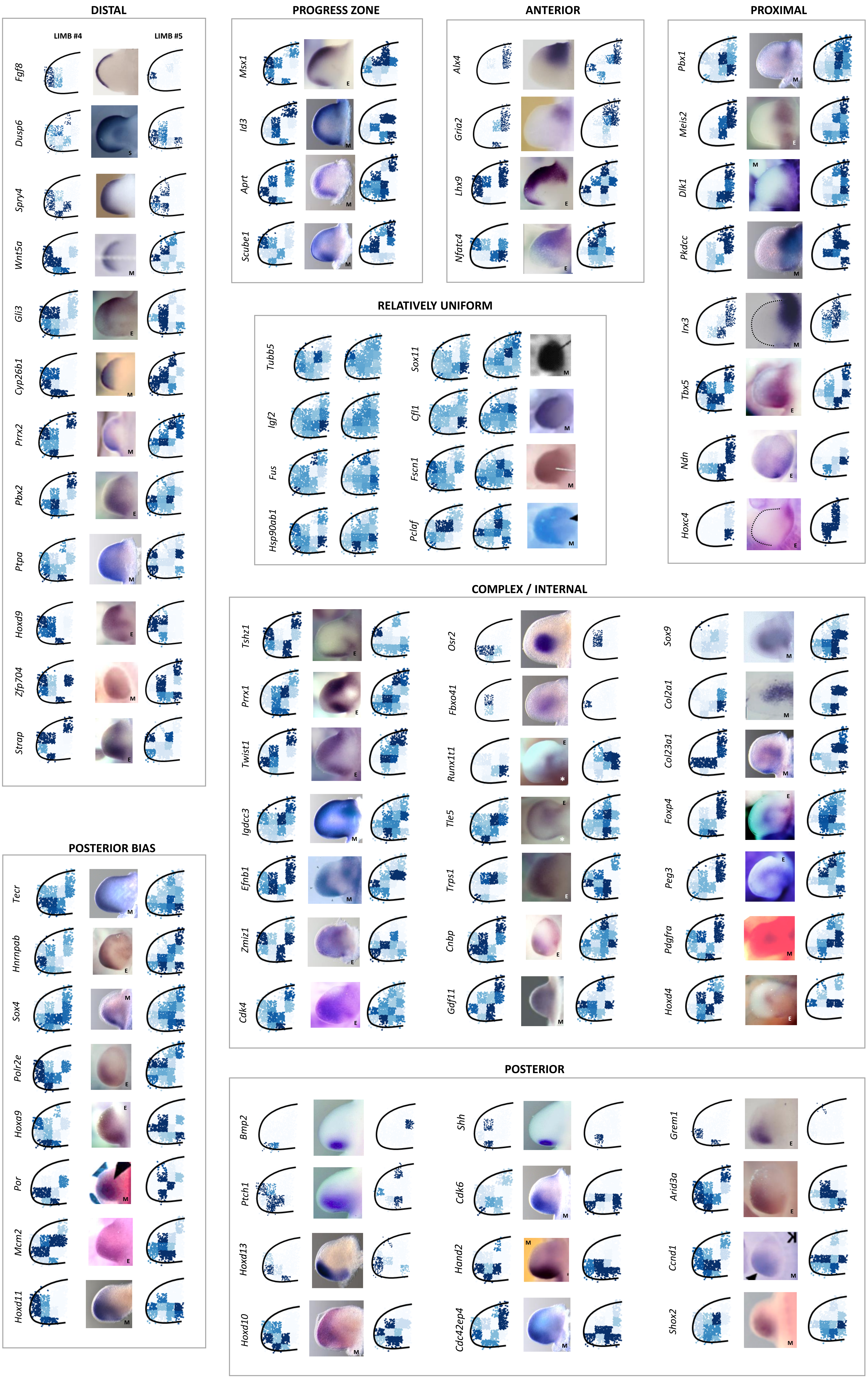
DISTAL
LIMB #4
LIMB #5
Fgf8
Dusp6
Spry4
Wnt5a
Gli3
Cyp26b1
Prrx2
Pbx2
Ptpa
Hoxd9
Zfp704
Strap
PROGRESS ZONE
Msx1
Id3
Aprt
Scube1
ANTERIOR
Alx4
Gria2
Lhx9
Nfatc4
PROXIMAL
Pbx1
Meis2
Dlk1
Pkdcc
Irx3
Tbx5
Ndn
Hoxc4
RELATIVELY UNIFORM
Tubb5
Igf2
Fus
Hsp90ab1
Sox11
Cfl1
Fscn1
Pclaf
COMPLEX / INTERNAL
Tshz1
Prrx1
Twist1
Igdcc3
Efnb1
Zmiz1
Cdk4
Osr2
Fbxo41
Runx1t1
Tle5
Trps1
Cnbp
Gdf11
Sox9
Col2a1
Col23a1
Foxp4
Peg3
Pdgfra
Hoxd4
POSTERIOR BIAS
Tecr
Hnrnpab
Sox4
Polr2e
Hoxa9
Por
Mcm2
Hoxd11
POSTERIOR
Bmp2
Ptch1
Hoxd13
Hoxd10
Shh
Cdk6
Hand2
Cdc42ep4
Grem1
Arid3a
Ccnd1
Shox2

**Figure S6: Unbiased scoring of C3PO gene expression reconstruction accuracy and precision.** The genes expression patterns for the 73 genes described above, and in the main results section. C3PO reconstructions for individual gene expression patterns are shown either side (limbs #4 and #5) of the measured gene expression pattern by WMISH (from the MGI GXD or Embrys databases). Blue is high expression and white is low gene expression as for Fig. 4 and S5. Genes are grouped together based on similar known gene expression patterns. We include four genes here for which we could not find expression patterns in the databases, because it is interesting to see how their relatively uniform patterns contrast dramatically with the “complex/internal” patterns in the section below. M = gene patterns found in the MGI database. E = gene patterns found in the Embrys database. S = Gene expression images generated by the Sharpe Laboratory. Remaining images come from publications listed in legend of Fig. 4. Asterisks (*) indicate two images of limb buds significantly older than limb #4 or #5. Here, we have used a generic outline for both limbs #4 and #5 to focus on the comparison between the gene expression patterns. Whole mount *in situ* hybridization (WMISH) patterns for *Osr2, Dlk1, Fbxo41, Pdkcc, Hoxd13, Cdk6* reprinted from[30] with permission from Elsevier. *Shox2* reprinted from[54] with permission from Elsevier, *Hand2, Alx4, Irx3, Hoxd11, Grem1, Cyp26b1, Fgf8* reprinted from[31] under a Creative Commons CC-BY-NC-ND licence, *Spry4, Gria2, Pbx1, Wnt5a* reprinted from[32] with a licence from Copyright Clearance Center on behalf of The Company of Biologists Ltd.*, Shh, Bmp2, Ptch1* reprinted from[33] under a Creative Commons licence, *Sox9* reprinted from[35] under a Creative Commons CC-BY licence, *Dusp6* generated by the Sharpe laboratory, *Gli3, Pbx2, Ptpa, Strap, Hnrnpab, Polr2e, Hoxa9, Mcm2, Msx1, Lhx9, Nfatc4, Meis2, Tbx5, Ndn, Hoxc4, Tshz1, Prrx1, Twist1, Zmiz1, Cdk4, Runx1t1, Tle5, Trps1, Cnbp, Foxp4, Peg3, Hoxd4, Grem1, Arid3a* reprinted from the Embrys database[34] with permission from Hiroshi Asahara, *Prx2* reprinted from[55] with a licence from Copyright Clearance Center on behalf of Elsevier Science & Technology Journals, *Ptpa, Zfp704, Tecr, Sox4, Id3, Aprt, Scube1, Igdcc3, Col23a1, Hoxd10, Cdc42ep4* reprinted from the MGI database with permission from Jordan Lewandowski, *Sox11* reprinted from[56] under a Creative Commons CC-BY-NC-ND licence, *Gdf11* reprinted from[57] under a Creative Commons CC-BY licence, *Cfl1* reprinted from[58] with a licence from Copyright Clearance Center on behalf of Elsevier Science & Technology Journals, *Fscn1* reprinted from[59] with a licence from Copyright Clearance Center on behalf of Elsevier Science & Technology Journals, *Pclaf* reprinted from[60] with a licence from Copyright Clearance Center on behalf of Elsevier Science & Technology Journals, *Efnb1* reprinted from[61] with a licence from Copyright Clearance Center on behalf of Elsevier Science & Technology Journals, *Col2a1* reprinted from[62] with a licence from Copyright Clearance Center on behalf of Elsevier Science & Technology Journals, *Pdgfra* reprinted from[63] with a licence from Copyright Clearance Center on behalf of Elsevier Science & Technology Journals. The *in situ* images for *Por* and *Ccnd1* were reprinted from[64,65] respectively, with a licence from Copyright Clearance Center on behalf of John Wiley & Sons - Books. Note that the *in situ* images for *Por* and *Ccnd1* may only be reused by contacting JOHN WILEY & SONS, INC. and requesting explicit permission. The links to the original articles[64,65] for these images are (figure 3d and figure 3k respectively for *Por* and *Ccnd1*):

https://anatomypubs.onlinelibrary.wiley.com/doi/full/10.1002/(SICI)1097-0177(199912)216:4/5%3C511::AID-DVDY19%3E3.0.CO;2-H

https://anatomypubs.onlinelibrary.wiley.com/doi/10.1002/dvdy.22598

### S7) Tissue dimension sizes for mapping fluorescence space to physical space.

The goal in this manuscript was to demonstrate C3PO's capability to accurately estimate relative cell positions and thus reconstruct the relative tissue configuration. Therefore, we simply adjusted the scales of the 3 axes to match the approximate proportions of the limb bud for limb buds #1-3 shown in Fig. 3. However, if desired a more exact scaling based on absolute physical space is possible by combining the FACS measured 3D point cloud data with the 3D image of the tissue measured predissociation.

This can be done by measuring the physical size of a pixel on the camera on the SPIM microscope and measuring the number of pixels spanning along each bleached axis of the tissue to produce a physical tissue size along each bleached axis. These physical sizes can be used to scale each dimension independently to convert FACS colour space to physical space so that the FACS measured point cloud (the cells) has the same relative dimension sizes of the physical tissue measured pre-dissociation in the SPIM microscope.

In the current manuscript the physical length of the main limb axes (proximal-distal, anterior-posterior and dorsal ventral) measured in this way directly correspond to the length of the axes bleached because we purposely bleached in parallel to them (Table S1). In 3D, we only performed this absolute scaling for the limb in Fig. S2 (top) since this was necessary to determine the absolute physical precision of C3PO (see section S8). In 2D (limbs #4 and #5), we simply scaled the raw FACS measured colour space to fit the outline of the limb bud measured from the predissociation limb image. This is effectively the same approach as described above but it is simpler to perform on 2D images than 3D images as it can be done in a basic graphics editor.

**Table S1:**

| | **Tissue axis sizes (in μm)** | | |
|---|---|---|---|
| | **Proximal-Distal** | **Anterior-Posterior** | **Dorsal-Ventral** |
| **Limb #1** | 762.5 | 851 | 496 |
| **Limb #2** | 1047 | 801 | 454 |
| **Limb #3** | 856 | 611 | 499 |
| **Limb Fig S2 top** | 726 | 770 | 428 |

**S8) C3PO spatial precision**

***a) Spatial precision theory***

To determine the spatial precision of the C3PO approach we first start with our signal (*y - active fluorophore concentration after bleaching*) over space (*x*) graph along each orthogonal axis of the 3D tissue. The active fluorophore concentration is normalised to the labelling efficiency control. This graph will inherently be noisy due to various features such as non-homogeneous staining, laser power fluctuations, background fluorescence, ambient light exciting the detector among others.

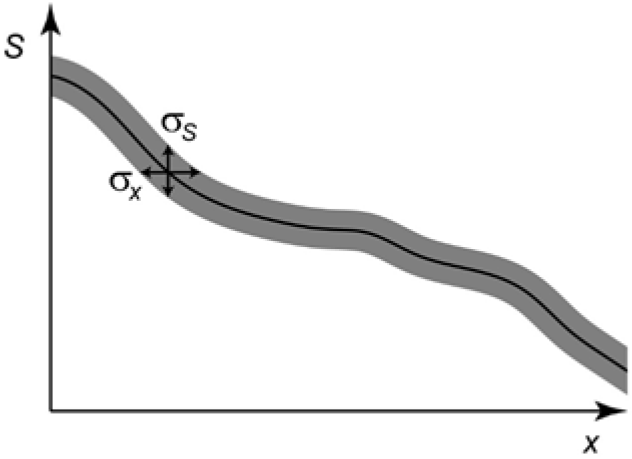


The noise (standard deviation) in the signal $\sigma_s$ can be converted into uncertainty (standard deviation) in the relative position of the cell σ along any given axis by the following equation:

$$\sigma_x = \frac{dx}{dS}\sigma_s \quad (1)$$

Assuming *S* is a linear function of *x*, this becomes

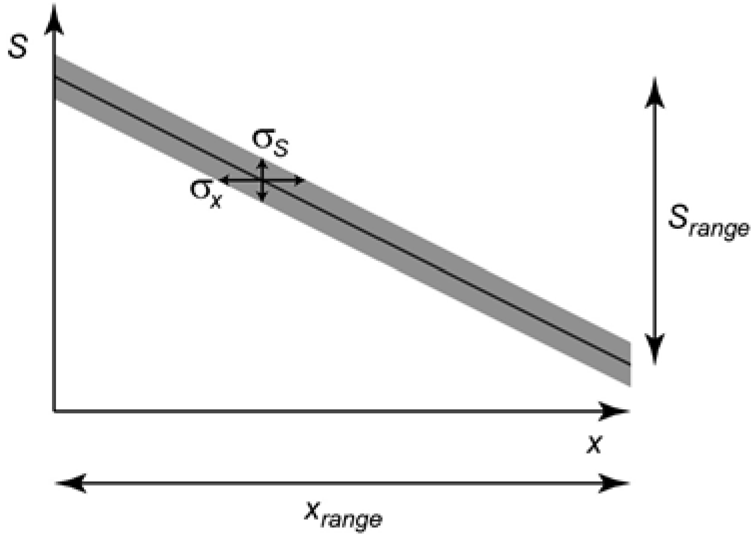

$$\sigma_x = \frac{x_{range}}{S_{range}} \sigma_s \qquad (2)$$

It is essential that the data used for measuring the spatial precision derives from the final FACS fluorescent signal measurements since these values encompass the cumulative error through the entire C3PO process (i.e. deriving spatial precision from the SPIM post bleaching pixel value image data would not take into account the error in reading the fluorescent signals of the dissociated cells). The spatial precision in the relative position of cells can be converted to spatial precision in absolute physical position independently for each dimension by assuming the FACS measured point cloud ($S_{range}$) is equivalent to the physical size of that bleached dimension as measured by the predissociation 3D SPIM image ($x_{range}$ that can be read from table S1).

It is useful to derive an extra concept when describing how well the C3PO system works, namely the number of resolvable segments per axis (NRSPA). The NRSPA can be determined by

$$NRSPA = \frac{x_{range}}{\sigma_x} = \frac{S_{range}}{\sigma_S} \qquad (3)$$

This concept relates to the utility of the approach to the biological community since it determines what questions are technically addressable via the C3PO approach.

#### b) *Spatial precision calculation for C3PO*

Using equation 2 we can state a spatial precision for each of the channels used in C3PO. In order to calculate the spatial precision we used dissociated limbs that were stained and bleached using the same parameters on the same Fortessa FACS analyzer. We measured $\sigma_s$ for each fluorophore from a stained but unbleached limb bud that had been dissociated with collagenase/dispase and analyzed on the Fortessa FACS Analyzer. The measured $\sigma_s$ were as follows:

**Table S2:**

| Normalized Fluorophore | $\sigma_s$ |
|---|---|
| Alexa405/TAMRA | 0.026 |
| 6FAM/TAMRA | 0.058 |
| Alexa647/TAMRA | 0.427 |

We then used a limb we had bleached in an all-or-nothing fashion for all 3 fluorophores since those fluorescent intensity levels after bleaching are equivalent to the maximum and minimum levels when we bleach a gradient and normalize the 3 channels we use for the *x, y* and *z* axes to the labelling efficiency control channel (TAMRA). These values (max-min) define the $y_{range}$ in the spatial precision calculation. The distance across each bleached axis (the $x_{range}$) can be calculated since we know the relationship between pixel and physical distance in our imaging system. The resulting values for a bleached limb are as follows:

**Table S3:**

| Normalized Fluorophore | $S_{range}$ (A.U.) | $x_{range}$ (bleached axis length in μm) |
|---|---|---|
| Alexa405/TAMRA | 0.23 | 428 (Limb Dorsal-Ventral axis) |
| 6FAM/TAMRA | 0.86 | 770 (Limb Anterior-posterior axis) |
| Alexa647/TAMRA | 6.09 | 726 (Limb Proximal-Distal axis) |

These numbers can then be plugged into equation 2 resulting in the following spatial precisions for our the mouse limb bud in our particular mesoscopic imaging system configuration:

**Table S4:**

| Normalized Fluorophore | Spatial precision (μm) |
|---|---|
| Alexa405/TAMRA | 48 |
| 6FAM/TAMRA | 52 |
| Alexa647/TAMRA | 51 |

Using equation 3 we determine the NRSPA as follows:
**Table S5:**

| Normalized Fluorophore | NRSPA |
|---|---|
| Alexa405/TAMRA | 8.8 |
| 6FAM/TAMRA | 14.8 |
| Alexa647/TAMRA | 14.3 |

#### c) Discussion on C3PO precision

From the theory, it can be seen very clearly that the precision of the C3PO system is influenced by three major factors

1) The noise in the signal: the smaller the noise $\sigma_S$ the better the spatial precision. Hence reducing factors including non-homogeneous staining, laser power fluctuations, background fluorescence, ambient light exciting the detector among others will improve the spatial precision of the C3PO approach.
2) The size of the tissue to which we are applying C3PO: the smaller the tissue the better the spatial precision.
3) The signal dynamic range $S_{range}$, since larger $S_{range}$ results in better precision (from eq. 2). Therefore, the precision can be improved by better staining or brighter

fluorophores and better ‘bleachability’ of the fluorophores (i.e. bleaching doesn’t saturate like it currently does for our fluorophores) to maximise the $S_{range}$.

Additionally another method to improve the precision in C3PO is by using multiple channels (as independent as possible) for each axis to increase the amount of spatial information that one can encode in the tissue. Note that the precision is not isometric for all axes in C3PO since each of these factors depends on the wavelength of light and opto-chemical characteristics of the fluorophore being used for each axis.

### Links to Creative Commons licences

***Creative commons licences:***
The CC-NY licence:
https://creativecommons.org/licenses/by/2.0/deed.en
The CC-NY-ND licence:
https://creativecommons.org/licenses/by-nd/2.0/deed.en
The CC-NY-NC-ND licence:
https://creativecommons.org/licenses/by-nc-nd/4.0/deed.en
